# Transposable elements drive regulatory and functional innovation of F-box genes

**DOI:** 10.1101/2024.12.17.628972

**Authors:** Miguel Vasconcelos Almeida, Zixin Li, Pedro Rebelo-Guiomar, Alexandra Dallaire, Lukáš Fiedler, Jonathan L. Price, Jan Sluka, Xiaodan Liu, Falk Butter, Christian Rödelsperger, Eric A. Miska

## Abstract

Protein domains of transposable elements (TEs) and viruses increase the protein diversity of host genomes by recombining with other protein domains. By screening 10 million eukaryotic proteins, we identified several domains that define multi-copy gene families and frequently co-occur with TE/viral domains. Among these, a Tc1/Mariner transposase helix-turn-helix (HTH) domain was captured by F-box genes in the *Caenorhabditis* genus, creating a new class of F-box genes. For specific members of this class, like *fbxa-215*, we found that the HTH domain is required for diverse processes including germ granule localisation, fertility, and thermotolerance. Furthermore, we provide evidence that HSF-1 mediates the transcriptional integration of *fbxa-215* into the heat-shock response by binding to Helitron TEs directly upstream of the *fbxa-215* locus. The interactome of HTH-bearing F-box factors suggests roles in post-translational regulation and proteostasis, consistent with established functions of F-box proteins. Based on AlphaFold2 multimer proteome-wide screens, we propose that the HTH domain may diversify the repertoire of protein substrates that F-box factors regulate post-translationally. We further demonstrate that F-box genes repeatedly and independently captured TE domains throughout eukaryotic evolution, and describe an additional instance in zebrafish. In conclusion, we identify recurrent TE domain captures by F-box genes in eukaryotes and provide insights into how these novel proteins are integrated within host gene regulatory networks.

## Introduction

In an influential analogy, François Jacob argued that evolution does not work like an engineer, striving for perfection in their creations (Jacob 1977). Instead, he asserted evolution acts as tinkerer “who does not know exactly what he is going to produce but uses whatever he finds around him”. In this sense, novelty is seldom created fully anew, *de novo*, but through the recombination of pre-existing material. He argued that “to create is to recombine” (Jacob 1977). These principles can be applied to protein evolution, with novel proteins arising by recombination of pre-existing modules. Several mechanisms can drive the recombination of protein modules, including exon shuffling, gene fusion, and transposition (Gilbert 1978; Patthy 1999; Babushok et al. 2007; Long et al. 2013).

Transposable elements (TEs) are mobile genetic elements detectable in most sequenced genomes, and often encode specialised protein machinery that is employed in their mobilisation (Wells and Feschotte 2020). There are many distinct types of TEs, categorised according to their sequence features, proteins encoded, and replication mechanism (Wicker et al. 2007; Bourque et al. 2018; Wells and Feschotte 2020). In brief, TEs are commonly divided in two broad classes based on the mechanism of mobilisation: class I TEs, or retrotransposons, transpose via an RNA intermediate, whereas class II TEs, or DNA transposons, employ a variety of mobilisation mechanisms exclusively via DNA intermediates (Wicker et al. 2007; Bourque et al. 2018; Wells and Feschotte 2020).

TEs are recognised as a major force in the evolution of eukaryotic genomes, driving innovation in a variety of ways (Bourque et al. 2018; Wells and Feschotte 2020; Almeida et al. 2022; Fueyo et al. 2022). TEs often contain and disperse transcription factor binding sites across eukaryotic genomes (Bourque et al. 2018; Almeida et al. 2022; Fueyo et al. 2022). This leads to their frequent repurposing as promoters or enhancers of endogenous genes, with the potential to establish or rewire cis-regulatory networks. The proteins encoded by TEs are also a source of innovation. They can contribute to the generation of novel proteins by recombining with pre-existing protein domains. For example, SETMAR proteins originated in primates via fusion of a Mariner DNA transposon with a SET histone methyltransferase (Robertson and Zumpano 1997; Cordaux et al. 2006). As another example in primates, CSB-PGBD3 emerged from a PiggyBac DNA transposon fusion with the Cockayne syndrome group B (CSB) gene (Newman et al. 2008). Larger-scale computational surveys of available genomic and proteomic data have identified additional proteins with co-occurring TE- and host-derived protein domains (Zdobnov et al. 2005; Cosby et al. 2021; Oggenfuss et al. 2024). One study, focusing on host-transposase fusion genes in tetrapod evolution, reported a tendency of TE-derived DNA-binding domains to fuse to host domains associated with the regulation of gene expression (Cosby et al. 2021). This study described KRABINER, a host-transposase fusion gene that binds DNA and regulates gene expression, akin to SETMAR and CSB-PGBD3 (Cordaux et al. 2006; Bailey et al. 2012; Gray et al. 2012; Tellier and Chalmers 2019; Cosby et al. 2021).

The increasing availability of sequenced genomes, along with their protein-coding and repetitive element annotations, provides a massive publicly available resource that can be leveraged to identify protein innovations. In this study, we screened 10 million eukaryotic proteins to identify biologically relevant TE- and virus-derived novelties. We describe two phylogenetically independent captures of TE domains by F-box genes in animals, and characterise in detail one instance in nematodes. In this case, one domain derived from Tc1/Mariner TEs was captured by an F-box gene and created a novel F-box gene family. These genes require the TE domain for diverse functions, including thermotolerance, fertility, and germ granule localisation. We further describe how Helitron TEs integrate one of these genes in a heat stress-responsive pathway.

## Results

### Multi-copy protein domains recurrently capture protein domains associated with TEs and viruses

We reasoned that protein domains usually in multiple copy in genomes, typically associated with multigene families, may retain fusions with TE- or virus-derived protein domains more often, given relaxed selective pressure on fusions which may be initially detrimental. This prompted our search for novel proteins with these domain architectures. We included protein domains associated with viruses in addition to TE domains, because long terminal repeat (LTR) retrotransposons are evolutionarily related to retroviruses, thus blurring the boundaries between these genetic elements (Eickbush and Malik 2007; Wells and Feschotte 2020).

To find instances of co-occurrence of TE- and virus-derived protein domains with multi-copy domains, we searched approximately 10 million unique eukaryotic proteins from the UniProt database (The UniProt Consortium 2023) and found 12,803 unique proteins in the major eukaryotic clades with such domain architectures (**Figure 1A** and **Table S1**). Of these, 9,471 have only one TE/viral domain, while 3,332 proteins display co-occurrence with multiple TE/viral domains, suggesting more complex fusions, possibly when several domains of one TE/virus are captured simultaneously (**Figure S1A**). There is a strong association between the size of the protein domain family and the co-occurrence with TE/virus-derived domains (**Figure S1B**). Reverse transcriptase, integrase, and peptidase domains are highly represented, associating with most multi-copy domains (**Figure 1B-C**). In turn, protein kinase, leucine-rich repeats, ankyrin repeats, GPCR, and F-box domains were amongst the multi-copy domains associated with most TE/viral domains (**Figure 1D**).

**Figure 1.**
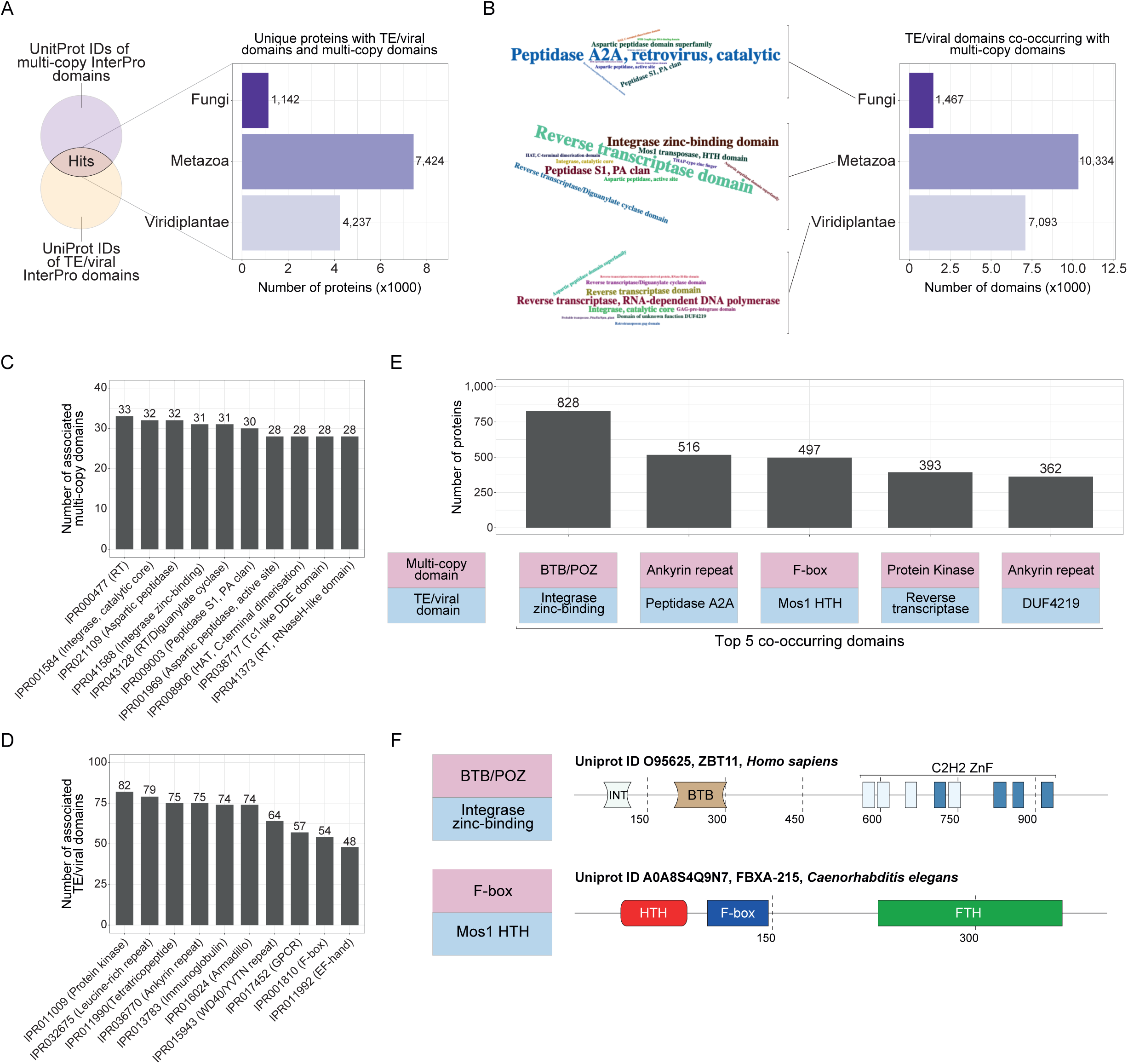
**Identification of eukaryotic proteins with TE/virus-derived and multi-copy domains**. (**A**) Simplified representation of the approach used to identify co-occurrence of multi-copy proteins domains with protein domains of TE/viral origin. Barplot depicts the number of unique proteins with protein domains of TE/viral origin in major eukaryotic clades. (**B**) Number of TE/viral protein domains co-occurring with multi-copy domains in major eukaryotic clades. Word clouds show the top 10 TE/viral domains in each clade. (**C**) The ten multi-copy domains with most associated TE/viral protein domains. (**D**) The ten TE/viral domains with most associated multi-copy domains. (**E**) Largest sets of proteins with the same co-occurring protein domains. (**F**) Protein domain architecture of representative proteins with co-occurring domains in the largest (BTB/POZ-Integrase zinc binding) and the third largest set of proteins (F-box-Mos1 HTH). FTH, FOG-2 homology domain; HTH, Helix-turn-helix; INT, integrase; RT, reverse transcriptase; Znf, zinc fingers.

To validate our approach, we searched for known instances of TE- or virus-derived domain fusions with multi-copy domains. The largest set of proteins with the same co-occurring domains, a BTB/POZ domain with an integrase zinc-binding domain (**Figures 1E-F, S1C** and **Table S1**), comprises a family of proteins in vertebrates with roles in neutrophil development by mediating repression of *TP53* (Keightley et al. 2017). The viral origin of the integrase-like domain was previously noted, and one of its amino acid residues responsible for the coordination of the metal ion is required for *TP53* repression (Keightley et al. 2017). Our approach also identified previously described SETMAR proteins (Robertson and Zumpano 1997; Cordaux et al. 2006; Tellier and Chalmers 2019) with SET and Mariner transposase domains (**Table S1**). Thus, our approach can identify known instances of co-occurrence of TE- or virus-derived domains with multi-copy domains.

### An F-box gene family with a TE-derived Helix-turn-helix domain in the *Caenorhabditis* genus

The third largest set of proteins with the same co-occurring domains have F-box (IPR001810) and Tc1/mariner Helix-turn-Helix (HTH) domains (IPR041426) structurally related to the N-terminal DNA-binding HTH domain of the Mos1 transposase of *Drosophila mauritiana* (**Figure 1E-F**)(Richardson et al. 2009). This large set of 497 proteins is phylogenetically restricted to the *Caenorhabditis* genus in nematodes (**Figure S1C**).

F-box domain-containing proteins adopt a variety of cellular functions, but are mostly known for their roles in the context of Skp, Cullin, F-box (SCF) E3 ubiquitin ligase complexes (Kipreos and Pagano 2000; Skaar et al. 2013). In specific, F-box proteins interact with protein substrates and bring them in close proximity to SCF complexes allowing substrate poly-ubiquitination and subsequent proteasomal degradation. A detailed study on gene families of ubiquitin-ligase adapters in *Caenorhabditis* has categorised *Caenorhabditis* F-box genes into 3 families: A1, A2, and B (Thomas 2006). Family A1 genes have an F-box and FTH domains, whereas family B genes have F-box and FBA2 domains (**Figure 2A** and **Table S1**). Family A2 was defined by having F-box and FTH domains plus an additional N-terminal domain, which was noted to be related to mariner transposases (**Figure 2A** and **Table S1**). The proteins we identified with co-occurring F-box and Tc1/mariner HTH domains (**Figure 1E-F**) correspond to the A2 family of F-box genes in the *Caenorhabditis* genus (**Figure 2A**). As the evolutionary history and functional roles of A2 F-box genes were not previously explored (Thomas 2006; Feschotte et al. 2009), we set out to do so.

**Figure 2.**
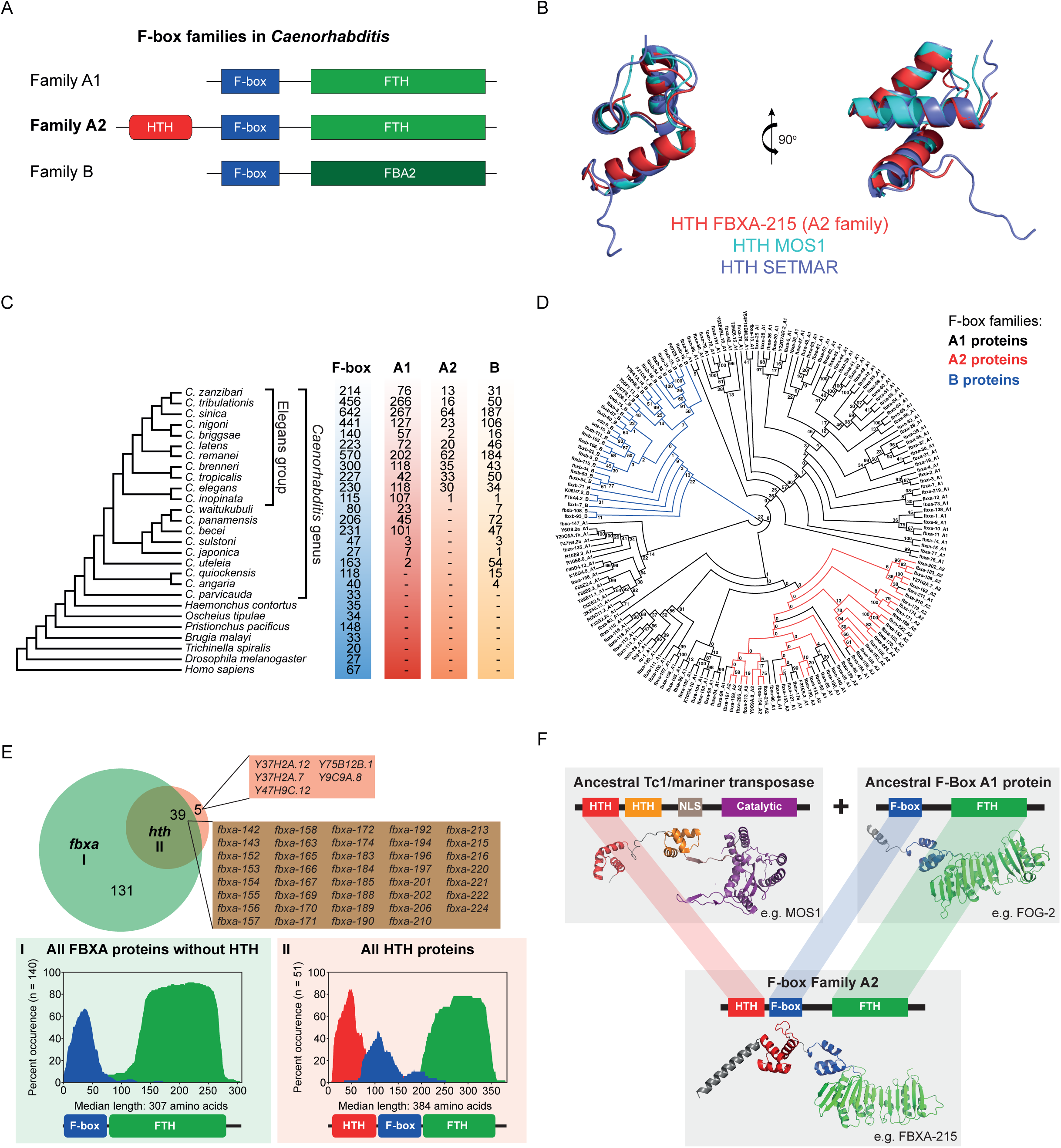
A subset of the F-box protein superfamily in *Caenorhabditis* nematodes has a Tc1/Mariner-derived Helix-turn-helix domain. (**A**) Schematics illustrating the domain architecture of proteins of the three F-box families in *Caenorhabditis*, as previously categorised (Thomas 2006). (**B**) Structural alignment of known TE-derived HTH domains of *Drosophila mauritiana*’s Mos1 (PDB: 3HOT) and *Homo sapiens*’ SETMAR (PDB: 7S03) with a representative HTH domain from FBXA-215, a protein of the F-box A2 family. All these HTH domains are of the simple tri-helical type. (**C**) Comparative genomic data from 27 species (nematodes, *Drosophila melanogaster*, and *Homo sapiens*) was used to date the origin of the A2 family. The cladogram was drawn based on previous phylogenomic analyses (Smythe et al. 2019; Stevens et al. 2019). The species belonging to the *Caenorhabditis* genus and the Elegans group are highlighted in the cladogram. For each species, the total number of F-box domain-containing genes is shown, as well as the members of all three families as in (A). (**D**) An alignment of all F-box proteins from *C. elegans* was used to construct a maximum-likelihood tree. The legend shows the colour code for all three gene families, see (A) for a schematic of their protein domain configurations. Numbers at internal branches indicate bootstrap support values (100 pseudoreplicates). (**E**) Venn diagram displaying the overlap between all the genes belonging to the *fbxa* class (including *F-box* A1 and A2 genes, indicated as I in the figure) according to Wormbase annotations, and all annotated *C. elegans* genes with a Tc1/mariner-derived HTH domain (IPR041426, indicated as II in the figure). The panels below depict the protein structure conservation for all the proteins encoded by the genes in groups I and II (all protein isoforms are included in the analysis). These graphs represent the percent of proteins with specific domains present along amino acid coordinates spanning the median protein length. Consensus protein structure is represented below. (**F**) Model for the emergence of the A2 family of F-box genes in the *Caenorhabditis* genus. In the ancestor of the Elegans group the N-terminal HTH domain of a Tc1/mariner transposase was captured on the N-terminus of an F-box gene of the A1 family, forming this novel F-box family, which subsequently expanded. Protein structures shown are AlphaFold predictions. FBA2; F-box associated domain 2; FTH, FOG-2 homology domain; HTH, Helix-turn-helix; NLS, nuclear localisation signal.

Given the ubiquity of HTH folds, we confirmed that the structure of the HTH domain of A2 family proteins is TE-derived and not related to other HTH folds. To do this, we performed structural alignments between proteins with TE-derived HTH folds of the same type, i.e. simple tri-helical type (Aravind et al. 2005). An excellent structural alignment is obtained with the HTH domains of A2 family proteins, SETMAR and Mos1 (**Figures 2B** and **S2A-C**, all-atom root mean square deviation, RMSD, between 0.7-2.1 Å). Conversely, alignment with non-TE-derived HTH folds of the simple tri-helical type is subpar (**Figure S2D-F**, RMSD between 2.8-6.4 Å). This supports a TE origin for the HTH domain of A2 family proteins.

Proteins with F-box domains are ubiquitous in eukaryotes, including animal genomes (**Figure 2C**). However, some nematode clades stand out with an expanded repertoire of F-box genes (**Figure 2C**) (Thomas 2006; Röseler et al. 2022). We revisited the composition of the F-box gene families in *Caenorhabditis* nematodes, following their initial categorisation (Thomas 2006). F-box genes with the domain structure of the A1 and B families of F-box genes are present only in the *Caenorhabditis* genus, suggesting these specific domain architectures are molecular innovations of this nematode genus (**Figure 2C**). Conversely, the A2 family is present only in the Elegans group within the *Caenorhabditis* genus, suggesting the HTH domain was captured by the F-box superfamily in the ancestor of the Elegans group, roughly 20 million years ago (Cutter 2008). It is relevant to note the three families only include a fraction of all *Caenorhabditis* F-box domain-containing genes. These observations suggest that a more comprehensive classification of the F-box superfamily using a broader taxonomic sampling might be needed. Phylogenetic analysis of *C. elegans* F-box proteins shows that B family genes form a monophyletic clade, while the A1 family has a patchier distribution (**Figures 2D and S2G**). Although the overall tree topology is not well supported, all the A2 family members fall into a subtree within the larger A1 family, suggesting a single origin for the A2 family (**Figure 2D**). Individual genes within the A2 subtree seem to have lost the HTH domain. Individual losses of other domains are also observed. For example, *fbxa-197* has lost the F-box domain and *Y37H2A.12* has no FTH domain. Altogether, these results support a single origin of the A2 family, followed by an expansion and subsequent losses of the HTH domain.

The majority of *C. elegans* genes with an annotated HTH domain (IPR041426) overlap with genes annotated as *fbxa* (39/44) and have an N-terminal HTH domain preceding F-box and FTH domains (**Figure 2E**, of note both A1 and A2 F-box gene families share the *fbxa* nomenclature). However, 5/44 HTH proteins are not annotated as *fbxa* and only 3/44 have isolated HTH domains (**Figures 2E** and **S3**). Further supporting a single origin for the A2 family, the HTH domain is present in F-box genes always N-terminally relative to F-box and FTH domains (**Figures 2E** and **S3**). Thus, the same overall domain architecture is maintained across the A2 family. All *C. elegans* HTH domain-containing proteins encode their HTH domains in one single exon (**Figure S3**). Therefore, the sequence of events following the capture is consistent with a previously proposed model (Cosby et al. 2021), involving TE insertion close to host transcripts, followed by exonisation of TE-derived protein coding sequences by alternative splicing.

Next, we interrogated the identity and origin of the captured TE. It is not clear whether the A2 ancestor received the HTH sequence directly from an endemic or horizontally transferred TE, or from an intermediate gene family that was the original recipient of the HTH domain. Further analysis of *Caenorhabditis* genes with HTH domains showed that this domain typically occurs either in combination with a transposase (IPR001888) or within an F-box context (either with an FTH, an F-box domain, or with both). These observations rule out the possibility of a secondary transition. Next, we searched for the most closely related sequences in species outside the Elegans group and in non-coding regions of the *C. elegans* genome. Phylogenetic analysis showed that one of the most closely related sequences corresponds to an intronic region that was annotated as a Mariner element in the *C. elegans* genome (**Figure S4**). This suggests that the HTH domain likely derived from an endemic TE. However, the overall tree topology is poorly supported, and we cannot completely rule out the possibility of an origin from a horizontally transferred TE.

Altogether, these results support a single origin for the A2 family within the *Caenorhabditis* genus, in the common ancestor of the Elegans group, when a Mariner HTH domain was captured by an F-box of the A1 family (**Figure 2F**).

### F-box A2 factors have signatures of purifying selection and a subset is expressed in the germline

Previous estimates of selection on F-box A1 genes identified stronger evidence of positive selection for the FTH domain, which interacts with substrates and positions them near the E3 ligase component of the SCF complex (Thomas 2006). In contrast, the F-box domain showed evidence of purifying selection, consistent with its role in binding to a Skp1 protein, thus connecting the F-box protein to the SCF complex (Thomas 2006). To acquire insights into the function and evolution of the A2 family, we analysed signatures of selection by examining the coding sequences of their protein domains (**Figure 2A**). This analysis supports a predominant signature of purifying selection for the F-box domain, and stronger evidence for amino acid residues under positive selection in the FTH domain (**Figure S5A**), in line with previous estimates (Thomas 2006). The HTH domain of F-box A2 proteins displayed a predominant signature of purifying selection (**Figure S5A**). In conclusion, the TE-derived HTH domain is predominantly evolving under purifying selection, which, in combination with the fact that it has been maintained in the genome for approximately 20 million years, strongly suggests it may be relevant for the functions of the F-box A2 gene family in the *Caenorhabditis* genus.

To further illuminate the potential functions of F-box factors, including the A2 family, we profiled their expression across development and in adult tissues of *C. elegans* using publicly available RNA-sequencing datasets (Ortiz et al. 2014; Almeida et al. 2019; Serizay et al. 2020). F-box genes are overall lowly expressed throughout development (**Figure S5B**), with the F-box B gene family showing higher expression in embryos. In contrast, the A1 family genes are overall more highly expressed than A2 family genes between L1 and L3 larval stages (**Figure S5B**). In terms of tissue-specificity in adult animals, more than 40% of A1 family genes are classified as having intestine-specific expression, whereas approximately 70% of B family genes are classified as lowly expressed in adults (**Figure S5C**). A2 family genes are the most versatile genes, with a similar proportion of genes being assigned as intestine-specific (26.6%), germline-specific (20.0%), and lowly expressed (23.3%, **Figure S5D**). The expression of A2 family genes in the germline led us to further explore expression of F-box genes in the germline. 19/116 A1 genes (16.4%), 12/30 A2 genes (40.0%), and 5/34 B genes (14.7%) had detectable expression in oogenic and/or spermatogenic gonads of *C. elegans* (**Figure S5D**). 15/19 germline-expressed A1 genes are expressed in both oogenic and spermatogenic gonads, while 4/5 B genes are expressed predominantly in oogenic gonads (**Figure S5E**). Again, A2 family genes display the most diverse expression patterns, with a similar proportion of genes predominantly expressed in gonads of each sex and in both sexes (**Figure S5E**).

In summary, the predominant signature of purifying selection on the TE-derived HTH domain of F-box A2 genes, along with their expression in the germline, suggest relevant functional roles in the germline.

### The HTH domain of F-box A2 family proteins is not involved in transcriptional regulation

A common denominator of the known roles of F-box proteins is the versatile mediation of protein-protein interactions, bridging different proteins or protein complexes (Kipreos and Pagano 2000; Skaar et al. 2013). Given the association of such a versatile protein module with a potentially DNA-binding HTH domain, we reasoned that F-box A2 proteins may have evolved a role in transcriptional regulation of TEs and/or endogenous genes, similar to other host-transposase fusion genes (Bailey et al. 2012; Gray et al. 2012; Tellier and Chalmers 2019; Cosby et al. 2021). As 40% of A2 genes are expressed in the germline (**Figure S5E**), such regulation could take place in germline tissues, which comprise a major stage of genetic conflict between TEs and their animal hosts (Ozata et al. 2019; Almeida et al. 2022). Among F-box A2 genes, *fbxa-192*, *fbxa-210*, and *fbxa-215* are the most highly expressed in the germline and in embryos, with *fbxa-215* being the most abundant (**Figure S5B, D**). These genes were selected to further interrogate the function of germline-expressed F-box A2 genes. We created mutant strains with complete or partial deletions of the coding sequence of these genes (**Figure S5F**). Furthermore, we endogenously tagged FBXA-192 and FBXA-215 with an N-terminal GFP.

Three lines of evidence argue against a transcriptional regulatory role of F-box A2 proteins. First, we profiled mRNA expression in embryos and young adults of wild-type N2 worms and *fbxa-215* mutants and did not find TE families differentially expressed between the two strains, in a consistent manner across developmental stages and growth conditions (**Figure S6A-D** and **Table S2**). Similarly, only a small number of protein-coding genes is differentially expressed in *fbxa-215* mutants and these changes are not consistent across developmental stages and growth conditions (**Figure S7A-F** and **Table S2**). Second, electrophoretic mobility shift assays failed to detect DNA binding of F-box A2 proteins and their HTH domains to the inverted repeat sequences derived from or similar to their ancestral Mariner TE (**Figure S8**). Third, FBXA-192 and FBXA-215 endogenously tagged with GFP display a dispersed localisation in the adult germline, with no nuclear localisation detected (**Figure S9A-B**).

We conclude that F-box A2 genes with high germline expression (FBXA-192/210/215) are unlikely to have a transcriptional regulatory role.

### F-box A2 family factors have roles in fertility and germline proteostasis

We further explored the localisation of F-box A2 proteins and observed GFP::FBXA-215, but not GFP::FBXA-192, localising to perinuclear germ granules in embryos (**Figures 3A, S9A, C**), as evaluated by co-localisation with DEPS-1, a factor known to localise to germ granules (Spike et al. 2008; Suen et al. 2020; Huang et al. 2024). Importantly, in-frame deletion of the HTH domain disperses GFP::FBXA-215(ΔHTH) from germ granules (**Figures 3A** and **S9D**), indicating the TE-derived HTH domain is required for the localisation of GFP::FBXA-215 to germ granules. In *fbxa-215* mutant animals germ granules look similar to wild-type, suggesting FBXA-215 is not required for germ granule condensation (**Figure S9E**).

**Figure 3.**
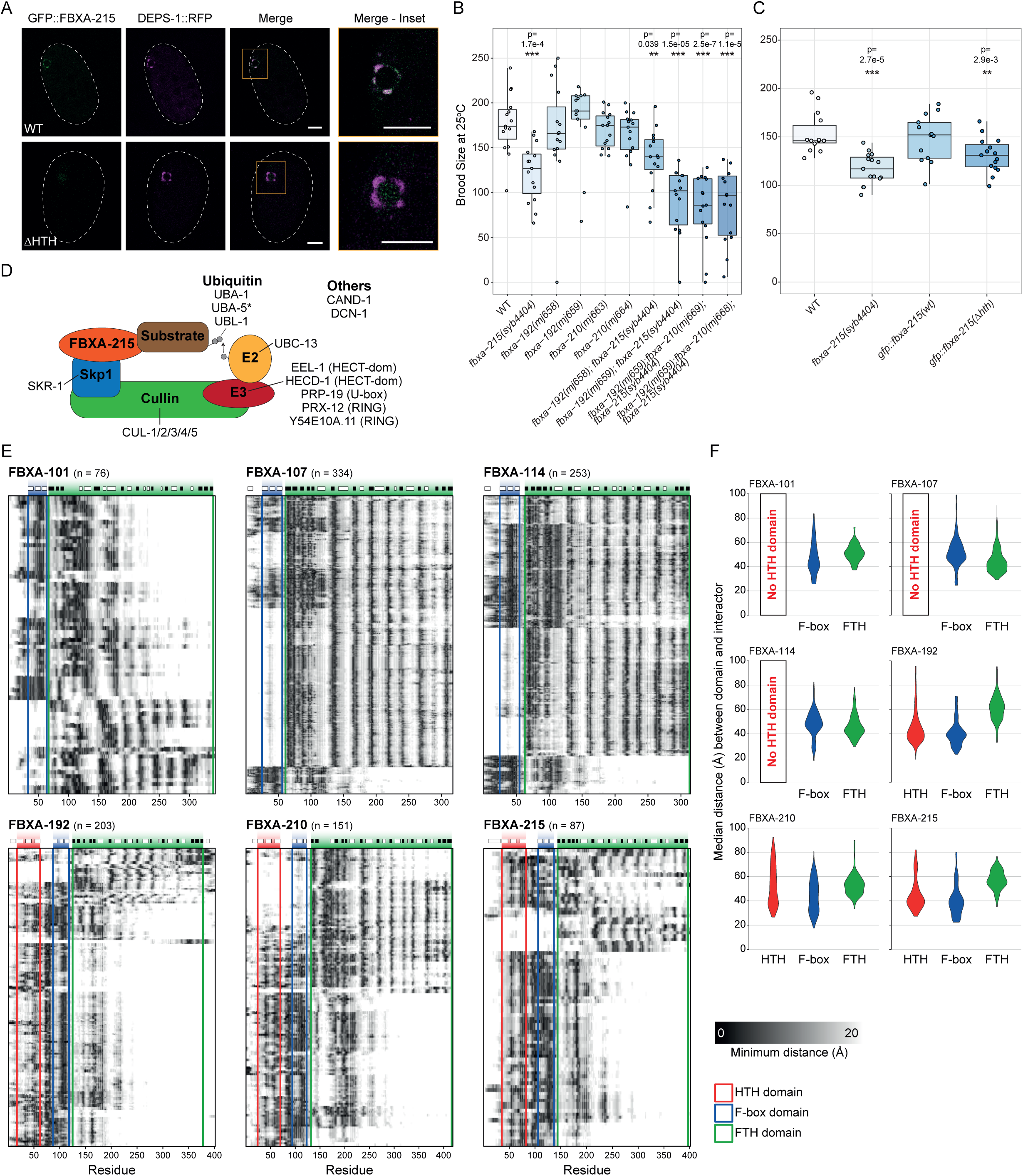
**Germline-enriched A2 F-box proteins in *C. elegans* are required for fertility and function in the context of SCF complexes**. (**A**) Confocal photomicrographs illustrating the localisation of GFP::FBXA-215 (green), DEPS-1::RFP (magenta), and their co-localisation in *C. elegans* embryos. Upper row of panels illustrate the localisation of these proteins in animals carrying a wild-type copy of GFP::FBXA-215. Conversely, the lower row of panels shows the localisation of these proteins in animals with a mutated GFP::FBXA-215, which lacks the TE-derived HTH domain. Insets focus on the P4 blastomere primordial germ cell (prior to its division into Z2 and Z3 primordial germ cells). Scale bars indicate 10 µm. Images represent single confocal planes images. Dashed lines represent the embryo outline. (**B-C**) Live progeny of the indicated strains at 25°C. Asterisks and p-values assessed by Mann–Whitney and Wilcoxon tests comparing wild-type (WT) N2 worms with the other strains. Horizontal lines represent the median, the bottom and top of the box represent the 25^th^ and 75^th^ percentile. Whiskers include data points that are less than 1.5 x interquartile range away from the 25^th^ and 75^th^ percentile. (**D**) Schematics representing the SCF complex factors detected in the interactome of FBXA-215. See detailed results in Table S3. (**E**) Heatmaps showing the minimum distance, in Å, between each amino acid residue of the F-box protein bait and the high-confidence interactors. Each row of the heatmap corresponds to one interactor. The domains of the F-box protein bait are indicated as coloured boxes, see colour key on the bottom right. The secondary structure of each F-box protein is shown on top of the respective heatmap, with the white and black boxes representing alpha helices and beta sheets, respectively. (**F**) Violin plots showing the median distance, in Å, between HTH, F-box, or FTH domains of F-box proteins to the high-confidence protein interactors obtained from the AlphaFold2 multimer screen.

Factors localising to germ granules are required for fertility (Kawasaki et al. 1998; Spike et al. 2008). Given the localisation of FBXA-215 to germ granules, we tested whether fertility is impacted by disruption of FBXA-215 and other germline-expressed F-box A2 genes. *fbxa-215* mutant animals display a mild fertility defect when compared to wild-type, whereas *fbxa-192* and *fbxa-210* show no fertility defects (**Figure 3B**). However, when these genes were mutated in combination with *fbxa-215*, fertility was more severely impacted, showing that F-box A2 genes contribute to fertility synergistically. *fbxa-215* mutants with an in-frame deletion of the TE-derived HTH domain have a fertility defect similar to the null allele, indicating that this domain is necessary for wild-type levels of fertility (**Figure 3C**).

To further understand the roles of F-box A2 factors in the germline, we explored their interactome. We expressed and purified F-box A2 constructs *in vitro* fused with a glutathione S-transferase (GST) solubility tag (overview of constructs in **Figure S10A**), incubated the purified proteins with *C. elegans* extracts and performed GST pulldowns followed by mass spectrometry. In line with the roles of F-box proteins in the context of SCF E3 ubiquitin ligase complexes, the interactome of FBXA-215 includes SCF complex factors (**Figure 3D** and **Table S3**). Two additional F-box A2 proteins, FBXA-192 and FBXA-210 also interact with SCF complex-associated proteins (**Figure S10B** and **Table S3**). Furthermore, gene ontology analysis of the interactomes of FBXA-215 and additional F-box A2 factors revealed enrichment for proteins related to the ribosome, mitochondria, and unfolded protein response (**Figure S10C** and **Table S3**), hinting at roles in stress responses.

Together, these data show that germline-expressed F-box A2 factors have roles in fertility, and that the TE-derived HTH domain of FBXA-215 is necessary to maintain normal fertility. Interactions with SCF complex factors support roles of F-box A2 proteins in proteostasis.

### The HTH domain may provide a protein-protein interaction platform to F-box A2 proteins

We performed GST pulldowns and mass spectrometry on additional germline-expressed F-box proteins, which do not encode an HTH domain (FBXA-101, FBXA-107, and FBXA-114). F-box proteins with an HTH domain tended to interact with a larger number of proteins compared to other F-box proteins (**Figure S10D**). In addition, we expressed the HTH domain and the C-terminal fragment of F-box A2 proteins separately (**Figure S10A**), and a pulldown-mass spectrometry revealed that HTH domains tended to interact with more proteins compared to C-terminal fragments (**Figure S10E-F**). These observations may suggest that the HTH domains of F-box A2 proteins mediate binding to diverse protein interactors, possibly expanding the post-translational regulatory capability of these F-box proteins.

To explore how the HTH domain may diversify protein-protein interactions of F-box proteins, we explored the protein-protein interfaces of F-box proteins. To do so, we performed AlphaFold2 multimer high-throughput screens using as baits germline-expressed F-box proteins with (FBXA-192, FBXA-210, and FBXA-215) or without (FBXA-101, FBXA-107, and FBXA-114) HTH domains, and modelled their interactions with the entire germline proteome. To attain a higher level of confidence in thresholding predictions, we first defined high-confidence interactors by analysing a set of true interactors and non-interactors with a benchmarking dataset. By applying these thresholds, we could define high-confidence interactors of F-box proteins (**Figure S11A**). SKR-1, the *C. elegans* ortholog of SKP1, emerged as a top interactor, with an observable interface with the F-box domain of all the F-box proteins tested (**Figure S11B**). This observation is in line with our mass spectrometry results (**Figure 3D**, and **Table S3**) and with the ability of F-box domains to interact with SKP1 proteins (Kipreos and Pagano 2000; Skaar et al. 2013). According to AlphaFold2 predictions, the high-confidence interactors of HTH-less F-box proteins make contacts preferentially with the C-terminal FTH domain, in line with the FTH domain being the domain responsible for recognition of protein substrates (**Figures 3E-F** and **S11C**). In F-box A2 proteins, most interactions occur in the N-terminus of these proteins, in the region overlapping the HTH domain (**Figures 3E-F** and **S11C**). These results agree with the mass spectrometry results (**Figure S10E-F**).

Overall, our data suggests that the TE-derived HTH domain provides a binding platform to recognise potential protein substrates, thus expanding the regulatory potential of F-box A2 proteins.

### Helitrons integrate F-box A2 genes into stress-responsive regulatory networks

Since results above pointed towards a role of F-box A2 proteins in the stress-response (**Figure S10C and Table S3**), we sought to understand how F-box genes are regulated and integrated with pathways responsive to stress in *Caenorhabditis*. In *C. elegans*, most F-box genes are located within two clusters on the arms of chromosomes II and V, in TE-rich regions (**Figure 4A-C**). Transcription factors can bind TE-derived sequences, regulating neighbouring genes, or integrating them into host regulatory networks, including networks responsive to stress (Horváth et al. 2017; Lanciano and Mirouze 2018; Almeida et al. 2022; Fueyo et al. 2022). We therefore developed an approach to address whether the genomic regions around F-box genes are significantly enriched in TEs. We observe variable enrichment of DNA, LTR, and Helitron elements in the vicinity of F-box genes in 11/17 *Caenorhabditis* genomes (**Figure S12**). We conclude that F-box genes tend to reside in variable TE-rich genomic regions, providing opportunity for new regulation to emerge.

**Figure 4.**
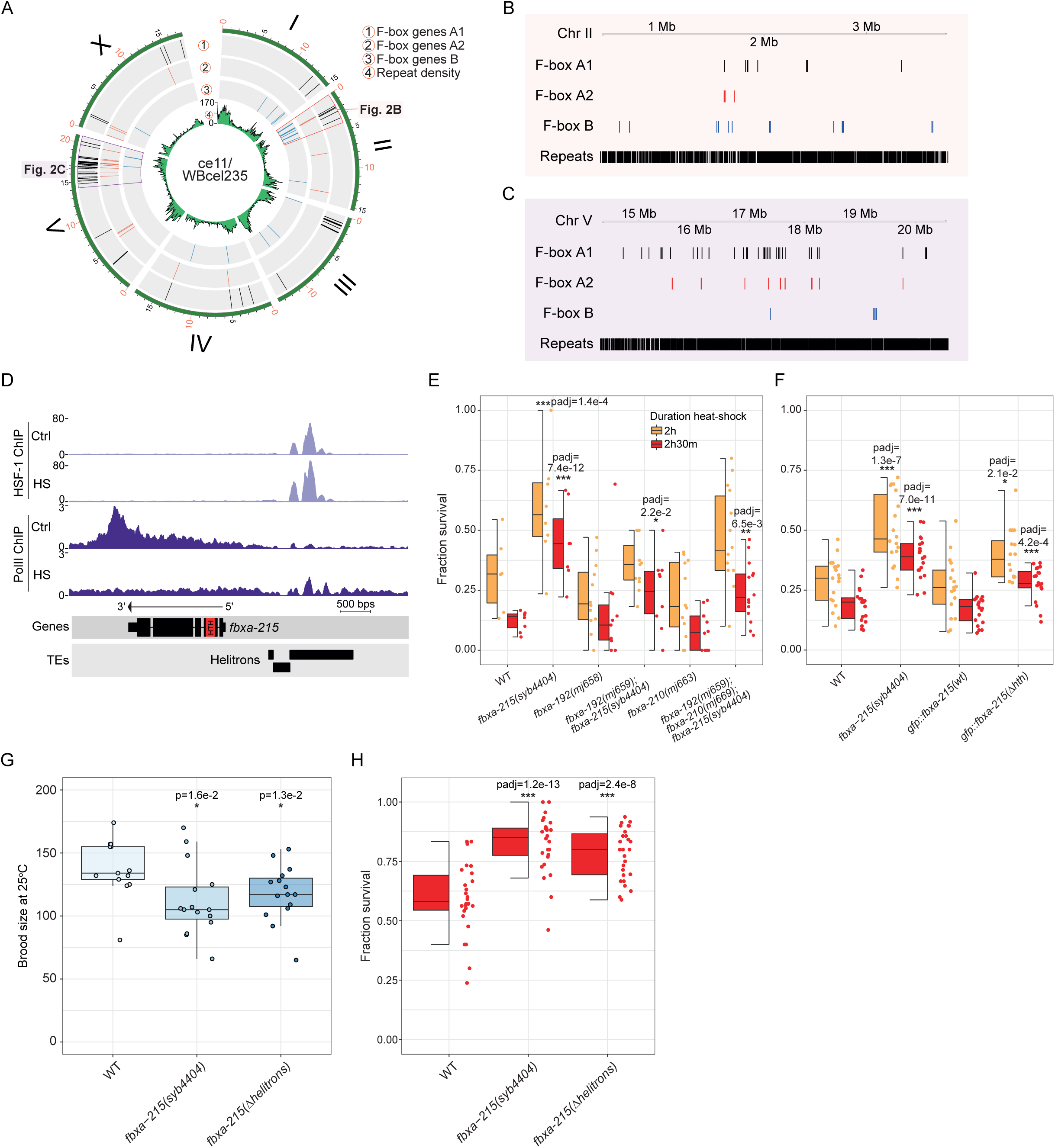
FBXA-215 modulates thermotolerance downstream of HSF-1-bound Helitrons. (**A**) Circos plot showing the location of all classes of F-box genes in the *C. elegans* genome. Inner track in green displays the repeat density across the genome. (**B-C**) Insets of some of the major clusters of F-box genes in chromosome II (B) and chromosome V (C) of *C. elegans*. (**D**) Genome tracks showing HSF-1 and RNA Polymerase II ChIP-sequencing data (Li et al. 2016) in the locus of *fbxa-215*. Ctrl, Control; HS, Heat-shock. The HTH domain is indicated in red in the *fbxa-215* annotation. (**E-F**) Worm survival 24 hours after 37°C heat-shock for 2h (orange) or 2h30m (red). Each figure represents two combined experiments. P-values show the results of Fisher’s exact tests comparing survival of worm strains versus survival of wild-type (WT) N2. (**G**) Live progeny of the indicated strains at 25°C. Asterisks and p-values assessed by Mann–Whitney and Wilcoxon tests comparing wild-type N2 worms with the other strains. (**H**) Worm survival 24 hours after 37°C heat-shock for 2h30m. These results represent three combined experiments. Asterisks and p-values denote the results of Fisher’s exact tests comparing survival of worm strains versus survival of wild-type N2. (E-H) Horizontal lines in the box represent the median, whereas the bottom and top of the box represent the 25^th^ and 75^th^ percentile. Whiskers include data points that are less than 1.5 x interquartile range away from the 25^th^ and 75^th^ percentile.

In *C. elegans*, we found Helitron TEs directly upstream of *fbxa-215* (**Figure 4D**). Helitrons are known to distribute binding sites for Heat-shock Factor 1 (HSF-1) in *Caenorhabditis* genomes (Garrigues et al. 2019; Schreiner et al. 2019). HSF1 is a conserved transcription factor in eukaryotes that coordinates transcriptional programmes in a variety of contexts, including development, metabolism, and in response to heat-stress (Vihervaara and Sistonen 2014; Gomez-Pastor et al. 2018). In *C. elegans*, HSF-1 has also been shown to regulate developmental programs and heat-stress responses (Chiang et al. 2012; Morton and Lamitina 2013; Brunquell et al. 2016; Li et al. 2016; Edwards et al. 2021). Previously published chromatin immunoprecipation sequencing (ChIP-seq) data (Li et al. 2016) support HSF-1 binding to the Helitrons upstream of *fbxa-215* (**Figure 4D**). Moreover, RNA Polymerase II elongation along *fbxa-215* is compromised upon heat-shock (**Figure 4D**), consistent with previously reported downregulation of *fbxa-215* after heat-shock (Brunquell et al. 2016). ChIP-seq and RNA-seq data from soma- and germline-specific depletion of HSF-1 (Edwards et al. 2021) further suggest that HSF-1 binding to the Helitrons near *fbxa-215* is mostly occurring in the germline (**Figure S13A-B**).

Next, to investigate the phenotypic effects of the regulation of *fbxa-215* by HSF-1, we quantified survival after heat-shock at 37°C for 2h and 2h30m. *fbxa-215* mutants displayed enhanced survival after heat-shock, when compared to wild-type N2 animals (**Figure 4E**). Other F-box A2 mutants did not display this phenotype and did not further enhance the phenotype of *fbxa-215* (**Figures 4E** and **S13C**). GFP::FBXA-215 did not change subcellular localisation upon heat-shock (**Figure S13D-E**). Of note, in-frame deletion of the HTH domain of FBXA-215 had increased survival compared to wild-type, phenocopying the null mutant and indicating the HTH domain affects thermotolerance (**Figure 4F**). Lastly, we determined the phenotypic impact of the Helitrons upstream of *fbxa-215*. To do so, we created mutants with the Helitrons deleted (not affecting the coding sequence of *fbxa-215*). In these mutant animals, under a non-stressful growth temperature of 20°C, *fbxa-215* is expressed at wild-type levels (**Figure S13F**). Nevertheless, disruption of the Helitrons upstream of *fbxa-215* affects fertility and thermotolerance, phenocopying *fbxa-215* null mutants (**Figure 4G-H**).

In summary, we report that F-box genes tend to reside in TE-rich regions across the *Caenorhabditis* genus. Also, Helitrons integrate *fbxa-215* into a thermal stress-responsive regulatory network in *C. elegans*.

### F-box proteins recurrently co-occur with TE protein domains

The tendency of F-box genes to occur in TE-rich regions provides opportunity for TE capture. This led us to interrogate whether F-box genes may have recurrently captured TE domains throughout eukaryotic evolution. Indeed, in our initial screen, we found 820 F-box domain-containing proteins in eukaryotes with potential fusions with domains associated with TEs or viruses (**Figure 5A-B** and **Table S1**). The majority of these unique proteins (492/820, 60%) correspond to the Mariner HTH domain capture in the *Caenorhabditis* genus that we identified and characterised above (**Figure 5B**).

**Figure 5.**
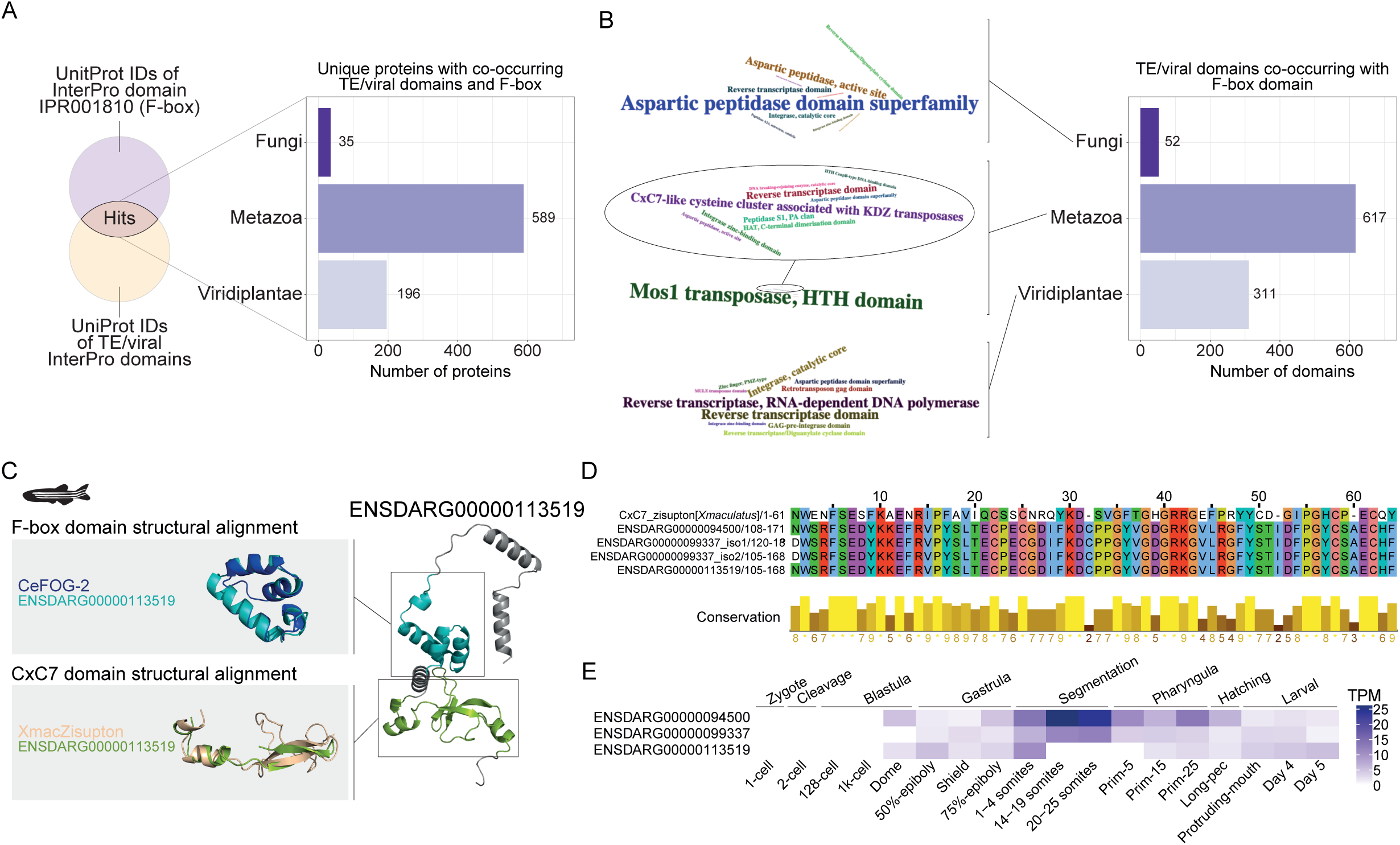
Additional example of a TE domain capture by F-box proteins in zebrafish. (**A**) Schematic depicting the approach to identify eukaryotic proteins with F-box domains (IPR001810) co-occurring with protein domains of TE/viral origin, a subset of the screen represented in Figure 1A. Barplot shows the number of unique F-box proteins proteins with TE/viral domains in major eukaryotic clades. (**B**) Number of TE/viral protein domains co-occurring with the F-box domain (IPR001810) in major eukaryotic clades. Word clouds show the top 10 TE/viral protein domains in each clade. In the metazoan word cloud, given the predominance of the Mos1 transposase HTH domain, mostly corresponding to the HTH capture described in Figure 2, we show an inset with the remaining terms. (**C**) Example of a protein with an F-box and a TE-derived CxC7-like cysteine cluster domain in zebrafish (*Danio rerio*). This is the protein product of ENSDARG00000113519 one of three zebrafish genes with these protein domains. Upper grey panel shows the structural alignment of the F-box domains of ENSDARG00000113519 and *C. elegans* FOG-2 (CeFOG-2). Lower grey panel shows the structural alignment of the CxC7-like cysteine cluster domain of ENSDARG00000113519 and *Xiphophorus maculatus*’ Zisupton. The zebrafish illustration was obtained from phylopic.org. (**D**) Multiple sequence alignment of the CxC7-like cysteine cluster domains of the Zisupton of *Xiphophorus maculatus (Xmaculatus*) and the three identified zebrafish genes (both protein isoforms encoded by ENSDARG00000099337 were included). (**E**) Expression of the F-box-KDZ genes identified, in transcripts per million (TPM), during zebrafish early development. Using data from White et al., 2017 (White et al. 2017).

Next, we sought to validate additional instances of F-box domain co-occurrence with TE or viral domains. We identified a set of F-box proteins in fishes with a zinc-chelating domain (CxC7-like cysteine cluster associated with KDZ transposases, IPR041300, **Figure 5C**) from KDZ (Kyakuja, Dileera, and Zisupton) TEs (Böhne et al. 2012; Iyer et al. 2014). The latter was hypothesised to be involved in the recognition of specific DNA sequences or chromatin proteins during transposition (Iyer et al. 2014). The CxC7 zinc-chelating domain in question (IPR041300) was defined based on the CxC7 zinc-chelating domain of Zisupton TEs in the platyfish *Xiphophorus maculatus* (Böhne et al. 2012). The zebrafish (*Danio rerio*) reference genome encodes three genes with F-box and CxC7 domains. The sequence and structure of their proteins confirmed the presence of an F-box domain, and of a TE-derived CxC7 domain (**Figure 5C-D**). Lastly, we used RNA-sequencing datasets of zebrafish embryogenesis and early developmental stages (Collins et al. 2012; White et al. 2017) to confirm the domain fusion and interrogate the expression of these genes. We observed reads spanning exons encoding the F-box and CxC7 domains, demonstrating that these two domains are transcribed and spliced together to encode a single polypeptide (**Figure S14**). The three F-box::CxC7 genes we identified are expressed in early zebrafish development (**Figure 5E**), with at least two of these showing prominent expression in segmentation. These expression patterns hint at developmental roles, which are yet to be explored.

These results suggest that F-box domain-containing proteins have recurrently captured TE- and/or virus-derived domains in eukaryotes.

## Discussion

In this work, by exploring a universe of 10 million unique proteins annotated in public databases, we identified over 12,000 proteins in eukaryotes with domains associated with TEs and viruses co-occurring with multi-copy protein domains, typically associated with multigene families (**Figures 1, S1** and **Table S1**). We hypothesised that TE/viral domains are more likely to be retained in gene families with functional redundancy, since the additional copies may buffer the impact of an initially deleterious chimeric transcript. Consistent with this, we observed a positive relationship between the number of proteins with a given domain and the number of instances of co-occurrence with TE/viral domains (**Figure S1B**).

TEs had a major role in eukaryotic genome evolution: 1) by providing their protein-coding sequences as building blocks that can be repurposed for other functions; and 2) by contributing to transcriptional regulation of endogenous genes, sometimes even wiring entire regulatory networks (Bourque et al. 2018; Almeida et al. 2022; Fueyo et al. 2022). We provide an example in *Caenorhabditis* nematodes that integrates these concepts, demonstrating both functional and regulatory contributions of TEs. We describe the capture of a Mariner HTH domain by a subset of the F-box superfamily (**Figure 2**), the F-box A2 family. We find striking that F-box A2 genes, which include a TE-derived protein domain with roles in diverse processes, are further integrated into host cis-regulatory networks by unrelated TEs, revealing multiple layers of TE-driven functional and regulatory novelty. For example, *fbxa-215* has Helitron TEs directly upstream of its coding sequence, which are bound by the transcription factor HSF-1 (**Figure 4D**). HSF-1 binding upstream of *fbxa-215* has phenotypic implications as deleting *fbxa-215*, its HTH domain, or the upstream Helitrons affects fertility and resistance to heat-stress (**Figure 4E-H**). These findings underscore the relevance of Helitron-driven genome evolution in the *Caenorhabditis* genus and beyond (Garrigues et al. 2019; Schreiner et al. 2019; Barro-Trastoy and Köhler 2024). These findings also exemplify how stress-responsive TE activity can integrate other genes in stress-responsive processes (Horváth et al. 2017; Lanciano and Mirouze 2018).

The F-box A2 gene family adds to a body of work in nematodes demonstrating that even organisms with a relatively low extant genomic proportion of TEs have a hidden abundance of TE-derived genomic novelties. For example, one recent study reported the discovery of TE-derived structural variation that modulates the expression of factors required for the biogenesis of particular small RNAs, and consequently affects the regulation of targets of these small RNAs (Zhang et al. 2024). Also, a germline cis-regulatory network was unveiled, wired by miniature inverted repeat TEs and a transcription factor related to a transposase (Carelli et al. 2022).

F-box A2 proteins and their HTH domains likely have a variety of biological roles. Here, we focused on three germline-expressed F-box A2 factors, out of a total of thirty A2 genes in *C. elegans*. We found FBXA-192/210/215 have synergistic roles in fertility, but they do not all contribute to thermotolerance (**Figure 3B-C**). This may be due to the fortuitous integration of FBXA-215, but not of FBXA-192/210, in the heat-stress response by Helitrons (**Figure 4D-F**). FBXA-215 thus seems to be pleiotropic, with potentially unrelated roles in fertility and thermotolerance. The more prominent role of FBXA-215 in fertility may be related to its localisation to germ granules, a localisation pattern not observed for FBXA-192 (**Figures 3A and S9A, C**). Other F-box proteins with roles in fertility have been described. FBXO24 is expressed in human and mice testes and is required for male fertility in mice (Kaneda et al. 2024; Li et al. 2024). FBXO24 was shown to interact with SCF complex factors and its role in fertility could be via SCF-driven post-translational regulation of target proteins (Kaneda et al. 2024; Li et al. 2024). FBXA-192/210/215 interact with SCF complex factors (**Figure 4D**), raising the possibility that post-translational regulation of specific protein targets underlies the fertility defect.

The proteome-wide prediction and modelling of F-box interactors suggests that the TE-derived HTH domain of F-box A2 proteins may alter protein-protein interactions, and diversify the repertoire of protein substrates SCF complexes can polyubiquitinate and help degrade (**Figure 3F-G**). The mechanistic basis should be further explored in subsequent studies. We propose that the co-option of TE-derived protein sequences as protein-protein interaction platforms contributing to post-translational regulation may represent a novel regulatory theme. Accordingly, we identified other domains known to mediate protein-protein interactions, such as BTB/POZ (Bonchuk et al. 2023), co-occurring with TE/viral domains (**Figure 1E-F** and **Table S1**).

We observed a similar proportion of A2 genes expressed in the germline and in the intestine (**Figure S5C**). FBXA-158 is an F-box A2 gene expressed in the intestine, which promotes thermotolerance in the context of the intracellular pathogen response targeting Microsporidia and viruses (Panek et al. 2020). The direct genomic vicinity of *fbxa-158* lacks Helitrons (**Figure S13G**), suggesting its role in thermotolerance emerged independently of the Helitron-HSF-1 axis in the germline. Other recent studies identified nematode F-box genes at the centre of hybrid incompatibility and toxin-antidote systems (Tikanova et al. 2024; Xie et al. 2024). Taken together, F-box genes are emerging as an evolutionarily labile toolkit that nematode genomes maintain and co-opt in diverse circumstances, with an impact in immunity, fertility, and viability. We propose that the variety of TEs enriched in the vicinity of F-box genes may enable their integration in various processes and in distinct cell types, in ways comparable to the integration of FBXA-215 in thermotolerance (**Figures 4A-D** and **S12**).

In addition to the F-box A2 family in *Caenorhabditis*, we report another class of proteins where an F-box domain co-occurs with a CxC7 zinc-chelating domain in fishes (**Figure 5**). This domain is defined by the CxC7 domain of platyfish Zisupton TEs (Böhne et al. 2012; Paysan-Lafosse et al. 2023). Of note, one of the protein isoforms of platyfish Zisupton TEs has been noted to include an F-box domain (Böhne et al. 2012), establishing precedence for the co-occurrence of F-box domains and Zisupton TE domains. The work described here, together with the previously described Zisupton-F-box domain association (Böhne et al. 2012) indicate recurrent association of F-box domains and TE domains. This may reflect recurrent captures of TE domains by F-box proteins, or, alternatively, capture of F-box domains by TEs. While we have not found any evidence supporting the latter scenario, we observed a tendency for F-box genes to locate in TE-rich regions of *Caenorhabditis* genomes (**Figures 4A-C** and **S12**). It will be useful to determine if this tendency extends beyond nematodes. If so, the residence of F-box genes close to an abundance of TEs may provide ample opportunities for captures throughout evolutionary time. The co-occurrence with TEs may be a general feature of gene families, not just of F-box genes, as gene families were recently shown to locate in TE-rich regions of eukaryotic genomes (Gozashti et al. 2024).

This study underscores the transformative role of TEs in eukaryotic evolution, contributing functional and regulatory novelties that influence complex traits, such as stress responses, immunity and fertility. Building on our findings and those of other screens (Cosby et al. 2021; Oggenfuss et al. 2024), future research should aim to uncover the full extent to which TE- and virus-derived domains diversify eukaryotic proteomes and generate functional novelty.

## Materials and Methods

### Identification of proteins with TE/viral and eukaryotic multi-copy protein domains

We first compiled and curated lists of InterPro protein domains (Paysan-Lafosse et al. 2023) associated with multi-copy genes and domains associated with TEs/viruses (**Table S1**). Next, we downloaded lists of Uniprot IDs and other metadata associated with these InterPro domains. We cross-referenced both sets of protein domains for shared UniProt IDs and concatenated all the IDs identified, along with relevant metadata (**Table S1**). The resulting data was imported to R (R Core Team 2021) to generate plots and conduct statistical tests. R scripts available at https://github.com/migueldvalmeida/F-box_TEs. We used the following R packages: tidyverse (Wickham et al. 2019), reshape2 (Wickham 2020), ggrepel (Slowikowski et al. 2023), wordcloud2 (Lang and Chien 2018), and taxonomizr (Sherrill-Mix 2023). Protein structure schematics on **Figure 1G** were created using DAVI (Saighi et al. 2021), with best match cascade. For BTB/POZ-integrase genes, we used the FASTA sequences of 160 mammalian ZBT11 proteins as DAVI input, while for F-box-Mos1 HTH we used 42 *C. elegans* proteins as input.

### Protein domain annotation and phylogenetic analysis

Comparative genomic data of several nematode species was downloaded from WormBase Parasite (version: WBPS16) (Howe et al. 2017). For *Pristionchus pacificus*, we used the latest gene annotation (version: El Paco gene annotation V3) (Athanasouli et al. 2020). Protein sequence for *Homo sapiens* and *Drosophila melanogaster* were downloaded from the Ensembl (release-93) and Ensembl Metazoa (release-40) ftp servers. In case of multiple isoforms per gene, the longest protein was chosen as the representative isoform. Protein domain annotation was done using the hmmsearch program of the HMMer package (version 3.3, hmmer.org, option -e 0.001) by searching against the Pfam database Pfam-A.hmm (version 3.1b2, February 2015) (Mistry et al. 2021). F-box containing genes were defined based on the presence of one of the domains PF00646 (F-box), PF12937 (F-box-like), PF13013 (F-box-like_2), PF15966 (F-box_4), PF18511 (F-box_5). The FTH and FBA2 domain were defined by the Pfam profiles PF01827 and PF07735, respectively. For phylogenetic analysis, F-box genes of all three families in *C. elegans* were aligned with the MUSCLE software (version 3.8.31) and the program raxml was run to compute a maximum-likelihood tree (version: 8.2.11, options: -m PROTGAMMAILG -f a) using 100 pseudoreplicates to compute bootstrap support values (Edgar 2004; Stamatakis 2014). To screen for homologous HTH domains, we performed a BLASTP search for *C. elegans* HTH domains against the NCBI nr database excluding any *Caenorhabditis* species and a BLASTP search against *Caenorhabditis* species outside of the Elegans group on WormBase ParaSite. Homologous non-coding sequences were identified by TBLASTN searches against the *C. elegans* genome (version: 2.10.1+, options: -max_target_seqs 2 -evalue 0.001). Phylogenetic analysis was performed as described above.

### Selection analysis

With the approach described in the last paragraph, we identified all the F-box A2 genes in the *Caenorhabditis* genomes and extracted their protein-coding and protein sequences from the respective transcriptomes and proteomes (downloaded from WormBase ParaSite, version WBPS16). The protein domain coordinates determined by HMMer were used to isolate the protein-coding and protein sequences of the domains from the full-length sequences. All the protein sequences of the HTH, F-box, and FTH domains were subsequently aligned using MAFFT v7.475 (Katoh and Standley 2013), using option --auto (selected FFT-NS-i strategy). Pal2nal v14 was used to create a reverse alignment from the protein sequence alignments and the respective protein-coding sequences (Suyama et al. 2006). The latter was used as input for selection tests in Datamonkey (Weaver et al. 2018), with Mixed Effects Model of Evolution (MEME) (Murrell et al. 2012). The results were plotted on an R framework (R Core Team 2021), using the following R packages: tidyverse (Wickham et al. 2019), reshape2 (Wickham 2020), patchwork (Pedersen 2023). The MEME method is sensitive to episodic selection and identified strong support for selection in 7/53 and 4/43 sites for the HTH and F-box domains (with p-value < 0.05), and 48/244 sites for the FTH domain.

### Visualisation of protein structure

Structural analysis and visualisation were conducted in Open-Source PyMOL v2.5 (The PyMOL Molecular Graphics System, v2.5, Schrödinger, LLC). All the structures were downloaded from AlphaFold Protein Structure Database and PDB. Structures from AlphaFold Protein Structure database: AF-O62474-F1 (*Caenorhabditis elegans* FBXA-215), AF-Q9XUX2-F1 (*C. elegans* FBXA-192), AF-E0R7K6-F1 (longest isoform of *C. elegans* FBXA-210), AF-G5EDX9 (*C. elegans* FBXA-101), AF-Q9U2P1 (*C. elegans* FBXA-107), AF-Q9U2X6 (*C. elegans* FBXA-114), AF-G5EBU7-F1 (*C. elegans* FOG-2), AF-G5EGQ5-F1 (*C. elegans* MBR-1), AF-P13528-F1 (*C. elegans* UNC-86), and AF-Q9GR61-F1 (*C. elegans* RBP-10). Structures from PDB: 3HOT (*Drosophila mauritiana* Mos1, obtained by X-ray diffraction) (Richardson et al. 2009), 7S03 (*Homo sapiens* SETMAR, obtained by X-ray diffraction) (Chen et al. 2022), and 2M3A (KNL-2, structure in solution obtained with NMR). For the structural alignments, only the helix-turn-helix domains were used, comprising the following amino acid ranges: FBXA-192 aa16-87, FBXA-210 aa24-76, FBXA-215 aa27-81, Mos1 aa5-55 (aligned only the first HTH), SETMAR aa330-394, MBR-1 aa147-193, UNC-86 aa363-437, RBP-10 aa1-67, and KNL-2 aa1-67.

The HTH, F-box, and FTH domains were annotated in the protein structures with the guidance of the domain coordinates from HMM as defined above, and according to extant annotations on Wormbase and Uniprot. Structural alignments of protein domains were done with Open-Source PyMOL v2.5 (The PyMOL Molecular Graphics System, v2.5, Schrödinger, LLC), using the align command. To compare the structural alignment of HTH domains we calculated all-atom RMSD (Å) values using PyMOL with the super command, cycles = 0. Results were then plotted on R, using packages tidyverse (Wickham et al. 2019) and reshape2 (Wickham 2020). ChimeraX (Goddard et al. 2018) was used to visualise the interaction between SKR-1 and F-box proteins.

The visualisation of the consensus protein structure of a group of proteins, as shown in **Figure 2E**, was produced using DomainViz (Schläpfer et al. 2021) via its web-server (https://uhrigprotools.biology.ualberta.ca/domainviz). We used default settings and the FASTA sequences of the groups of proteins indicated in **Figure 2E** as input.

As there were no structures available for fish F-box::CxC7 and for *Xiphophorus maculatus* Zisupton protein (UniProt ID: G3KL21, used only the CxC7 domain, residues 1333-1393 according to InterPro annotation), we used AlphaFold Colab (Jumper et al. 2021) to model their structures (monomer model, run_relax). Amino-acid sequences and predictions available at https://github.com/migueldvalmeida/F-box_TEs.

### Caenorhabditis elegans genetics and culture

Animals were cultured on HB101 bacteria according to standard laboratory conditions (Brenner 1974). *C. elegans* were grown at 20°C unless otherwise specified. We used the Bristol strain N2 as the reference wild-type strain. **Table S4** lists all strains used in this study. Strain design and other analysis were aided by Wormbase resources (Davis et al. 2022).

### CRISPR/Cas9 genome editing

Mutants were generated using CRISPR/Cas9 genome editing. Cas9 ribonucleoprotein complexes were pre-assembled *in vitro* and injected into the germlines of wild-type N2 animals, or *fbxa-192(mj659); fbxa-215(syb4404)* double mutants (strain SX3717, see **Table S4**).

Desalted and deprotected 2’-O-methylated Edit-R guide RNAs (gRNAs) were used (Horizon Discovery Biosciences). These gRNAs are able to bind an Edit-R synthetic tracrRNA (Horizon Discovery Biosciences) *in vitro*, due to complementary sequences on their 3’ end (Jinek et al. 2012). See **Table S5** for a list of gRNAs used. The injection mixture comprised 25 mM KCl, 7.5 mM HEPES-KOH pH = 8.0, 1 μg/μL tracrRNA, 160-200 ng/μL of each gRNA, and 80-160 ng/μL *dpy-10* gRNA (co-CRISPR, see below). In some experiments, 5 ng/μL pCFJ104 body wall muscle mCherry reporter was also added to the mix. These components were mixed by gentle tapping, incubated at 95°C for 5 minutes and allowed to cool down at room temperature (for at least 5 minutes). Then, recombinant Cas9 (CP02, PNA Bio) was added to a final concentration of 250 ng/μL, mixed and incubated for 5 minutes at room temperature. Finally, 20 ng/μL of a DNA *dpy-10* repair template was added along with nuclease-free water for a total volume of 10 μL. The mixture was then spun at 21,000 x G between 2-5 minutes on a table-top centrifuge. Worms were immobilised on 2% agarose microinjection pads, in Halocarbon oil 700 (Sigma-Aldrich). Injection was performed with InjectMan 4 (Eppendorf) and Eppendorf femtotips using an Olympus IX71 microscope. F1 worms were selected based on the dumpy co-CRISPR phenotype (Kim et al. 2014), due to *dpy-10* mutation, and were screened by PCR and Sanger sequencing to identify and confirm deletion alleles. Additional strains were created by SunyBiotech, using CRISPR/Cas9 genome editing (see **Table S4**). CRISPR-Cas9 mutants used in the experiments shown in this work were outcrossed 4x with wild-type N2 animals.

### Brood size determination

Done as previously described (Almeida et al. 2018; Dietz et al. 2021). In short, 15 L3 worms of each strain (grown at 20°C) were individually isolated and grown at 25°C. After onset of egg laying, animals were transferred to a new plate every day, until no eggs were laid for two consecutive days. Live progeny was counted approximately 24 hours after removing the parent. Worms that died before egg laying terminated, for example, by dehydration on the side of plate, were excluded from the analysis. Each experiment was performed at least two independent times. Statistical significance was tested using two-sided Wilcoxon-Mann-Whitney tests.

### Thermotolerance experiments

We performed the following protocol based on previous studies (Panek et al. 2020) and published experimental recommendations (Zevian and Yanowitz 2014). Two gravid adults of each strain were isolated into a fresh plate, allowed to lay eggs for approximately 1 hour, then removed off the plate. Eggs laid during this 1-hour period were allowed to develop at 20°C for 48 hours producing a synchronized population of L4 stage animals. After 48h, plates were sealed with parafilm and the L4 worms were heat-shocked for 120 or 150 minutes at 37°C. After heat-shock plates were moved to room temperature for 30 minutes, and then to a 20°C incubator. Worms were scored for survival approximately 24h after heat-shock ended. Worms were scored as dead when they failed to respond to touch and did not show any pharyngeal pumping. Each experiment was performed at least two independent times. Statistical significance was tested using Fisher’s exact test.

### *In vitro* GST fusion protein expression and GST pulldowns

*Expression of GST fusions in E. coli*. Constructs were cloned, expressed and purified as previously reported (Almeida et al. 2018). Plasmids encoding GST-F-box fusion proteins were retransformed in Rosetta 2(DE3) Singles Competent Cells (Novagen, 71400-3), as per the manufacturer’s instructions. One colony was inoculated in a 5 mL pre-culture, supplemented with Ampicillin and Chloramphenicol (100 µg/ml and 25 µg/ml, respectively), and grown overnight at 37°C. This pre-culture was used to inoculate a 200-250 mL culture of LB, supplemented with Ampicillin and Chloramphenicol (100 µg/mL and 25 µg/mL, respectively) and grown at 37°C, up to an OD600 of 0.5-0.9. Then, protein expression was induced with 1 mM isopropyl β-D-thiogalactoside (IPTG, Melford, I56000) and incubated overnight at 18°C. Next, bacteria were collected by centrifugation (4°C, 4000rpm, 15-30 minutes) and pellets were frozen at -80°C.

*GST-on bead purification*. Bacteria pellets were resuspended in Lysis Buffer (50 mM Tris pH 7.5, 150 mM NaCl, 1 mM DTT, 0.1% Triton-X, and protease inhibitors, cOmplete Mini, EDTA-free, Roche, #4693159001). Next, the samples were sonicated with a Sonics Vibra-Cell instrument 4 times for 2 minutes (amplitude = 30%), with 5-10 minute pauses between cycles. Debris were pelleted by centrifugation (4°C, 4000rpm, 30-45 minutes) and discarded, supernatant was filtered through a 0,20 µM Sartorius Minisart filter. A 250-300 µL slurry of Glutathione Sepharose™ High Performance beads (GE Healthcare, 17527901) was washed 3 times with 1 mL lysis buffer. Centrifugation steps in washes were conducted at 800 x G for 3 minutes. The cell lysate was then incubated with the beads between 2-3 hours at 4°C with end-to-end mixing. After incubation, beads were washed between 10-15 times with wash buffer (50 mM Tris pH 7.5, 150 mM NaCl, 1 mM DTT, and protease inhibitors, cOmplete Mini, EDTA-free, Roche, #4693159001). and, after the last wash, suspended 1:1 in wash buffer. 5% glycerol was added to the proteins for short-term storage at 4°C. To examine the purity of the preparation, 1x LDS buffer (Life Technologies, #NP0007) was added to the samples, and samples were subsequently loaded onto a denaturing NuPAGE Bis-Tris 4-12% gel (Life Technologies, #NP0335BOX) in a NuPAGE MOPS SDS running buffer (Life Technologies, #NP0001) at a constant voltage of 150 V. Gels were stained with InstantBlue Coomassie stain (Abcam, #ab119211) and imaged.

*Worm extract preparations*. A synchronised population of young adult worms was obtained by bleaching gravid adults grown at 20°C to obtain embryos, allowing the embryos to hatch overnight in M9, with gentle mixing, bringing the synchronised L1s on plate, and growing them at 20°C. These synchronised animals were collected and lysed in Lysis Buffer (25 mM Tris HCl pH 7.5, 150 mM NaCl, 1.5 mM MgCl2, 1 mM DTT, 0.1% Triton X-100 and protease inhibitors: cOmplete Mini, EDTA-free, Roche, #4693159001). Lysis was performed by sonication (10-20 cycles of 30 seconds ON and 30 seconds OFF) in a Bioruptor Plus (Diagenode). For embryo collection, large populations of gravid adults grown at 20°C were bleached, embryos were thoroughly washed with M9 buffer, and frozen in Lysis Buffer using liquid N2. Lysis was performed by grinding frozen embryo pellets and douncing with 40 strokes, piston B. Lastly, lysates were cleared by centrifugation (15 minutes at 21,000 x G, 4°C) and protein concentration was measured using Bradford Protein assay according to manufacturer’s instructions (Bio-Rad, #5000006).

*GST pulldowns*. 5 µg of GST-F-box beads were washed 3 times with pulldown wash buffer (25 mM Tris HCl pH 7.5, 150 mM NaCl, 1.5 mM MgCl2, 1 mM DTT and protease inhibitors: cOmplete Mini, EDTA-free, Roche, #4693159001). Subsequently, 300-700 µg of worm extract (either from young adult animals or embryos) was incubated with the beads for 3 hours, at 4°C, with end-to-end mixing. Beads were washed 3 times (with pulldown wash buffer) and resuspended in a total 25 µL volume including 1x LDS buffer (Life Technologies, #NP0007) and 100 mM DTT. Lastly, beads were boiled at 95°C for 10 minutes and frozen at -20°C until processing of samples for mass spectrometry took place (see below).

### Mass spectrometry

*In-gel digest*. In-gel digestion followed previously established procedures (Shevchenko et al. 2007). Samples underwent electrophoresis on a 4–12% Bis-Tris gel (NuPAGE, Thermo Scientific, #NP0321) for 8 minutes at 180 V in 1x MES buffer (Thermo Scientific, NP0002). Each lane was excised and cut into approximately 1 mm x 1 mm pieces using a sterile scalpel and transferred into one well of a 96 well hydrophilic low protein binding filter plate (Merck Millipore, MSBVN1210). Gel pieces were destained using destaining buffer (50% 50 mM ammoniumbicarbonate buffer, or ABC, pH 8.0, 50% ethanol ) at 37°C with vigorous agitation until completely destained. Subsequently, gel pieces were dehydrated by immersing them in 100% acetonitrile for 10 minutes at 25 °C with shaking. The gel pieces were incubated in reduction buffer (50 mM ABC, 10 mM DTT) at 56°C for 60 minutes. The reduction was followed by incubation in alkylation buffer (50 mM ABC, 50 mM iodoacetamide) for 45 minutes at room temperature in the dark. Gel pieces were washed with digestion buffer (50 mM ABC) for 20 minutes at 25°C. Then, gel pieces were dehydrated again by incubation in pure acetonitrile until gel pieces got white and hard. Samples were further dried at 80°C until the filter membrane of the 96 well plate turned white. The dried gel pieces were rehydrated in a trypsin solution (50 mM ABC, 1 µg MS-grade trypsin per sample, Serva Electrophoresis 37286) and the filter plate placed on top of a 96 deep well collection plate (Eppendorf, 951032603). Gel pieces were incubated overnight at 37°C. The plate assembly was centrifuged at 300 x G for 2 minutes. Flowthrough containing tryptic peptides was collected and combined with additional elutes obtained by treating the gel pieces with extraction buffer (50 mM ABC, 30% acetonitrile) twice and a further step with pure acetonitrile for 10 minutes at 25°C, shaking at 300 rpm. The dehydration step with acetonitrile was repeated until gel pieces got white and hard. The sample containing tryptic peptides was reduced to 10% of the original volume using a Concentrator Plus (Eppendorf, #5305000304, settings V-AQ) to remove acetonitrile and purified using the StageTip protocol.

*Stage tip purification*. Stage tip purification followed mainly a previously described method (Rappsilber et al. 2007). Desalting tips were created by stacking two layers of C18 material (Affinisep AttractSPE, #SPE-Disk-Bio-C18-100-47.T1.20) within a 200 µl pipet tip. These tips were primed using pure methanol. Following activation, they underwent successive rinses with solution B (80% acetonitrile, 0.1% formic acid) and solution A (0.1% formic acid) each for 5 minutes before application of the tryptic peptide samples. Subsequently, a final wash with solution A was performed. When utilized, peptides were eluted using solution B. The resultant samples were centrifuged in a Concentrator Plus for 10 minutes to remove acetonitrile and were adjusted to a volume of 14 µl using solution A.

*MS analysis*. For MS analysis a volume of 5 µL desalted and eluted peptides of each sample were injected and separated on a nanocapilary column (New Objective, 25 cm long, 75 µm inner diameter) packed in-house with C18 (Dr. Maisch GmbH) for reverse-phase chromatography. This setup was interfaced to an EASY-nLC 1000 system (Thermo Scientific) coupled to a Q Exactive Plus mass spectrometer (Thermo Scientific). Peptides were eluted from the column employing an optimized 2-hour gradient ranging from 2 to 40% of 80% MS grade acetonitrile/0.1% formic acid solution at a flow rate of 225 nL*min-1. The mass spectrometer was operated in a data-dependent acquisition mode, conducting one MS full scan followed by up to ten MS/MS scans using HCD fragmentation. All raw files were processed using MaxQuant (version 2.4.2.0)(Cox and Mann 2008) and matched against the *C. elegans* Wormbase protein database (version WS269, 60000 gene transcripts) and the Ensembl Bacteria *E. coli* REL606 database (version from September 2018, ASM1798v1, 4533 gene transcripts) for proteins originating from the *E. coli* feeding strain. Carbamidomethylation (Cys) was set as a fixed modification, while methionine oxidation and protein N-acetylation were considered as variable modifications. Enzyme specificity was set to trypsin with a maximum of two miscleavages. LFQ quantification without fast LFQ was performed, requiring at least 2 LFQ ratio counts, and the match between run option was activated. Filtering and analysis were done in R (R Core Team 2021). Interactors of GST::F-box fusion proteins were defined by significant enrichment over GST control. For GST pulldowns from young adult extracts, interactors were defined by fold change > 2 and p-value < 0.1. For GST pulldowns from embryo extracts, interactors were defined by fold change > 1.5 and p-value < 0.1. Gene ontology was conducted on sets of F-box protein interactors using g:Profiler web server’s g:GOST functional profiling with the *Caenorhabditis elegans* genome and default parameters, including driver terms (Kolberg et al. 2023). See detailed mass spectrometry and gene ontology results in **Table S3**.

### Electrophoretic mobility shift assay (EMSA)

After purification (see above), GST fusion proteins to use in EMSA were eluted from the beads 5 times with 150 μL of elution buffer (wash buffer with 10 mM reduced glutathione, pH readjusted to 7.5 with NaOH). The eluates were combined, and the sample concentrated and buffer-exchanged back to wash buffer on a 3k Amicon Ultra-0.5 mL centrifugal filter (Merck Millipore). Protein concentration was quantified by measuring OD595 in a Bradford Protein Assay (BioRad, #5000006) relative to bovine serum albumin standards. 5 μg of the sample was run on a denaturing gel to examine the purity of the preparation and correct protein size.

PAGE-purified single-stranded DNA oligonucleotides (**Table S5**) were annealed by incubation in a heating block set at 89°C and allowed to cool down to room temperature overnight. The annealing reaction comprised 5 μL of each 100 μM complementary oligonucleotide, 2 μL annealing buffer (200 mM Tris-HCl pH 7.5, 100 mM MgCl2, 1M KCl) and 8 μL nuclease-free water. Binding reactions were carried out in wash buffer (50 mM Tris-HCl pH 7.5, 150 mM NaCl, 1 mM DTT, EDTA-free protease inhibitors Roche, 2.5 % glycerol) in a total volume of 15 μL with purified protein and 77.3 nM DNA. After incubating at 25°C for 15 minutes, 2.5 μL of loading dye (50 mM Tris-HCl pH 7.5, 150 mM NaCl, 33 % glycerol, 0.1 % bromophenol blue) was added. The samples were run on a native 1.5 % UltraPure agarose (Invitrogen) gel in 1X TBE pH = 7.5 for 80 minutes (4°C, 90 V constant voltage). The gel was stained with 1X SYBR Gold (Invitrogen) in 1X TBE pH = 7.5 for 40 minutes rocking, and then imaged.

### Microscopy

A flat 2% agarose pad was prepared on a glass slide. To prepare gravid adult animals for imaging, animals were picked into a droplet of M9 to wash off the excess *E. coli* bacteria the worms grow on. Subsequently, animals were transferred to another droplet of M9 on the agarose pad, supplemented with 2.5-5% sodium azide for immobilisation. Embryos *in utero*, or extruded during the immobilisation process were also imaged. This approach was employed to create the images in **Figure S9**, using a Nikon Ti2-E inverted microscope (NIS-Elements AR image acquisition software, v5.42.06) equipped with a Plan Apo VC 20x/0.75 air objective, a Plan Apo VC 60x/1.2 water objective, a Nikon DS-Qi2 camera, green (Nikon, #MXR00704) and red (Nikon, #MXR00711) fluorescence filters, as well as bright field. The images in **Figure 3A** were created by imaging embryos, which were obtained through bleaching gravid adults, followed by thorough washes in M9 buffer. Afterwards, embryos were introduced into an M9 droplet on a 2% agarose pad and immediately imaged on a Leica SP8 confocal on a DM6000 upright microscope (Leica LAS X image acquisition software, v3.5.7.23225), equipped with an HC PL APO CS2 100x/1.4 oil objective. Imaging was done with sequential scanning with detection ranges maximised for signal. Images shown in **Figure 3A** represent a single confocal plane taken with a 1AU pinhole at 580 nm. Images were processed using Fiji v2.14 (Schindelin et al. 2012), by adjusting brightness and contrast.

### RNA extraction and mRNA sequencing

N2 wild-type and *fbxa-215(syb4404)* animals grown at 20°C were synchronized by bleaching and overnight hatching in M9 buffer with gentle mixing. Synchronized L1 animals were brought onto HB101-seeded NGM plates the next day and grown at 20°C or at 25°C. Synchronised young adult worms were collected approximately 36 hours (batch grown at 25°C), or 48 hours (batch grown at 20°C) after plating. Animals were washed off plates with M9, further washed with M9, washed one last time in nuclease-free water, and frozen in dry ice. To collect embryos, synchronized gravid adult worms were bleached approximately 47 hours (batch grown at 25°C), or 56 hours (batch grown at 20°C) after plating. After 2 washes in ice-cold M9 and a final wash in nuclease-free water, embryos were frozen in dry ice.

A previously published RNA extraction protocol was used with minor alterations (Almeida et al. 2019). Frozen worm aliquots were thawed, 300 μL of TRIzol (Life Technologies, 15596026) were added and mixed vigorously. Six freeze-thaw cycles were employed to dissolve animals, specifically by freezing tubes in liquid nitrogen for approximately 30 seconds, followed by thawing for approximately 3 minutes at 37°C. After each cycle, tubes were mixed vigorously. After the sixth freeze-thaw cycle, samples were spun down at 20,000 x G for 3 minutes and the supernatant was subsequently collected into a fresh tube. Next, 1 volume of 100% ethanol was added to the sample and mixed vigorously. Finally, the mixtures were loaded onto Direct-zol columns (Zymo Research, R2060) and manufacturer’s instructions were followed (in-column DNase I treatment was included). Quality control of samples, library preparation (non-directional, with poly-A enrichment), and mRNA sequencing (Illumina, PE150) was performed by Novogene.

### Bioinformatic analysis

*mRNA-sequencing analysis of datasets generated in our study*. Illumina adapters and reads with low-quality calls were filtered out using Trimmomatic v0.39 (Bolger et al. 2014) with options SLIDINGWINDOW:4:28 MINLEN:36. Quality of raw and trimmed fastq files was assessed with fastQC v0.11.9 (https://www.bioinformatics.babraham.ac.uk/projects/fastqc/) and summarised with multiQC v1.11 (Ewels et al. 2016). Gene expression was quantified from trimmed reads using salmon v1.5.1 (Patro et al. 2017), with options --seqBias --gcBias --validateMappings -l A. DESeq2 (Love et al. 2014) and custom scripts (available at https://github.com/migueldvalmeida/F-box_TEs) were used to calculate normalised and TPM counts, generate plots and conduct statistical tests on an R framework (R Core Team 2021). See R packages used below, in the end of this section.

Trimmed fastq files were mapped to the *C. elegans* genome (WBcel235) using HISAT2 v2.2.1 (Pertea et al. 2016). Resulting SAM files were converted to BAM format and sorted with samtools v1.10 (Li et al. 2009): 1) samtools view -bS; 2) samtools sort; and 3) samtools index. To create bigwig files, the BAM alignment files were used as input to bamCoverage v3.5.1, part of the deepTools package (Ramírez et al. 2016), using options --normalizeUsing CPM -of bigwig --binSize 10. Bigwig files of biological replicates were combined using wiggletools (Zerbino et al. 2014) mean and wigToBigWig v4 (Kent et al. 2010). Genome tracks were plotted with custom scripts (available at https://github.com/migueldvalmeida/F-box_TEs) using the Gviz (Hahne and Ivanek 2016) and GenomicFeatures (Lawrence et al. 2013) packages on an R framework (R Core Team 2021).

To quantify TE expression at the TE family level, we first mapped trimmed reads to the *C. elegans* genome (ce10) using STAR v2.5.4b (Dobin et al. 2013) with options --readFilesCommand zcat --outSAMtype BAM SortedByCoordinate --outFilterType BySJout --outFilterMultimapNmax 150 --winAnchorMultimapNmax 150 --alignSJoverhangMin 8 --alignSJDBoverhangMin 3 --outFilterMismatchNmax 999 --outFilterMismatchNoverReadLmax 0.04 --alignIntronMin 20 --alignIntronMax 10000000 --alignMatesGapMax 100000000. The resulting BAM files were used as inputs for TEtranscripts v2.2.1 (Jin et al. 2015) with options --stranded no --SortByPos.

As TEtranscripts input we used ce10 gene and TE annotations. The latter was downloaded from the list of TEtranscripts-compatible annotations created by the Hammell laboratory (available here: https://hammelllab.labsites.cshl.edu/software/). DESeq2 (Love et al. 2014) and custom scripts (available at https://github.com/migueldvalmeida/F-box_TEs) were used to calculate normalised counts, generate plots, and conduct statistical tests on an R framework (R Core Team 2021).

For the analysis above, the following R packages were used: tidyverse (Wickham et al. 2019), lattice (Sarkar et al. 2008), eulerr (Larsson et al. 2022), genefilter (Gentleman et al. 2023), pheatmap (Kolde 2019), reshape2 (Wickham 2020), ggrepel (Slowikowski et al. 2023), biomaRt (Durinck et al. 2023), tximport (Love et al. 2023), RColorBrewer (Neuwirth 2022), ashr (Stephens et al. 2023), ggpubr (Kassambara 2023), GenomicFeatures (Lawrence et al. 2013), patchwork (Pedersen 2023), and UpSetR (Conway et al. 2017).

*Analysis of publicly available datasets*. Publicly available ChIP-sequencing processed data (Li et al. 2016; Edwards et al. 2021) was used to generate genome tracks. Bedgraph files were downloaded from GEO (accessions GSE81521 and GSE162063) and converted to bigwig files with bedGraphToBigWig v2.8 (Kent et al. 2010). BigWigs were used to generate genome tracks with custom scripts (available at https://github.com/migueldvalmeida/F-box_TEs) using the Gviz (Hahne and Ivanek 2016) and GenomicFeatures (Lawrence et al. 2013) packages on an R framework (R Core Team 2021).

Publicly available raw RNA-sequencing data from *C. elegans* (Almeida et al. 2019; Edwards et al. 2021) was downloaded from public repositories (GEO, accession number GSE162064 and SRA, accession number PRJNA497368) and analysed exactly as described above for the data generated in this study (except for TEtranscripts analysis, which was not conducted). In addition, we used publicly available raw RNA-sequencing data from zebrafish (Collins et al. 2012; White et al. 2017), downloaded from public repositories (BioProject numbers PRJEB2333 and PRJEB12982). This data was analysed exactly as described above (TEtranscripts analysis was not done), except that reads were mapped to the Ensembl zebrafish genome (Danio rerio GRCz11, or GCA_000002035.4).

Lastly, we used publicly available quantitative (tables of normalised counts) and qualitative (categorisation of genes according to tissue-specific expression) data (Ortiz et al. 2014; Serizay et al. 2020).

### RT-qPCR

RNA was extracted from synchronized young adult animal populations as described above. Subsequently, reverse transcription (RT) and quantitative PCR (qPCR) were conducted as previously described (Almeida et al. 2018), with minor changes. RT was performed with 700 ng of RNA using ProtoScript First Strand cDNA Synthesis Kit (New England Biolabs, #E6300), according to manufacturer’s instructions and using an oligo d(T)23 VN. After preparation, RT was diluted 1:3 times in nuclease-free water. Afterwards, qPCR reactions were prepared using Power SYBR Green Master Mix (Thermo Fisher Scientific, #4367659) according to manufacturer’s instructions. Reactions were prepared for a total of 10 µL, including a primer final concentration of 300 nM, and 1 µL of template RT. A StepOnePlus Real-Time PCR System (Thermo Fischer Scientific) was used to run the qPCR reactions. Cycling conditions were the following: Standard run; 5 minutes at 95°C; [40 cycles of 95°C for 15 seconds and 60°C for 45 seconds]; melt curve calculation [15 seconds at 95°C, 1 minute at 60°C and 15 seconds at 95°C (1.6°C/s of increment in temperature)]. Technical duplicates and biological quadruplicates were used. Analysis was performed using the ΔΔCT method (Schmittgen and Livak 2008). *pmp-3* was used as a normalisation factor (Hoogewijs et al. 2008). Error bars represent the standard deviation of the four biological replicates. Primers used are indicated in **Table S5**.

### Proteome-wide structural interaction screen

Receiver operating characteristic (ROC) analysis of ColabFold (Jumper et al. 2021; Mirdita et al. 2022) predictions was performed with a library of 255 pairs of true interactors, including entries from *Caenorhabditis elegans*, curated from Vreven et al., 2015 (Vreven et al. 2015), Chen et al. 2003 (Chen et al. 2003), CASP15 (https://predictioncenter.org/casp15), PDB (https://www.rcsb.org), and 490 pairs of true non-interactors curated from Blohm et al. 2014 (Blohm et al. 2014). Prediction performance was evaluated by assessing the area under the curve of the ROC curve for several parameters, with pTM and ipTM selected for downstream thresholding using the Kolmogorov-Smirnov statistic.

The library of preys was compiled based on a previous study (Ortiz et al. 2014), using information of *C. elegans* genes known to be expressed in spermatogenic or oogenic gonads. Entries with duplicated sequences were removed from the proteome/prey library and matched with the sequence of each entry in the FBXA/bait library in preparation for MSA search and 3D model prediction (without Amber relaxation) with ColabFold. Information on pTM, ipTM and atomic positions was extracted from each prediction to assess interaction potential and for analysis in R (R Core Team 2021). Atoms separated by a distance smaller than the sum of their van der Waals radii were considered clashed and not used for analysis.

### Association between TEs and gene families

TEs were annotated in *Caenorhabditis* species listed in **Table S4** using EDTA v2.0.0 with the option --sensitive 0 (Ou et al. 2019), and unclassified TE copies were filtered out. Gene annotations and PFAM domain annotations were downloaded from WormBase Parasite (Howe et al. 2016; Howe et al. 2017). TE enrichment by PFAM was calculated using a set of custom scripts (available at https://github.com/migueldvalmeida/F-box_TEs). Briefly, for each species, up to ten classified TE copies closest to every gene +/- 500 bp were counted, and counts per gene were then summed for each PFAM domain. A thousand random distributions of TE counts per PFAM were generated by a bootstrapped shuffling of TE positions across all scaffolds. An empirical cumulative distribution function was used to calculate the distance of the observed number of TEs per PFAM from the random distributions, and p-values were corrected for multiple comparisons to generate scores.

### Localisation of *C. elegans* F-box genes and TEs

Ensembl BioMart (Harrison et al. 2024) was used to collect the genome localisations of all F-box A1, A2, and B genes (identity of these genes can be found in **Table S1**). A circos plot was created with Circos v0.69-8 (Krzywinski et al. 2009) to visualise F-box gene localisation together with TE distribution. TE Density tracks are displayed on the circos plot as the number of features per 100 kilobase.

Genome tracks with zoomed-in genomic regions on chromosomes II and V were plotted with custom scripts (available at https://github.com/migueldvalmeida/F-box_TEs) using the Gviz (Hahne and Ivanek 2016) and GenomicFeatures (Lawrence et al. 2013) packages on an R framework (R Core Team 2021).

## Data Availability

RNA sequencing data generated in this study have been deposited to GEO under accession number GSE251877. Proteomics data are available at the ProteomeXchange Consortium via PRIDE under the accession number PXD048290. All other data sets used in this manuscript are publicly available on WormBase, WormBase ParaSite, Genbank, PDB, GEO, SRA, UniProt, and InterPro.

## Acknowledgements

We are grateful to all members of the Miska and Rödelsperger labs, as well as Ben Luisi, for discussion and suggestions. We would like to thank Honour McCann, and Oliver Weichenrieder for helpful comments. We thank Nicola Lawrence of the Gurdon Institute Imaging Facility for microscopy training and assistance. We are grateful to Juan Carlos Rueda Silva, David Jordan, and Bryan Tan for technical assistance, and to Marc Ridyard for excellent laboratory management. We are indebted to the Gurdon Institute Media Kitchen for providing reagents and media. Some strains were provided by the CGC, which is funded by NIH Office of Research Infrastructure Programs (P40 OD010440). We acknowledge SunyBiotech for generating some of the worm strains used in this study.

## Author contributions

Conceptualisation: M.V.A., C.R.; Investigation: M.V.A., Z.L., P.R.-G., L.F., J.S., C.R.; Data Curation: M.V.A., Z.L., P.R.-G., C.R.; Formal Analysis: M.V.A., Z.L., P.R.-G., A.D., L.F., J.L.P., J.S., X.L., C.R.; Visualisation: M.V.A., C.R.; Writing – Original Draft: M.V.A., C.R.; Writing – Review & Editing: all authors contributed; Project Administration: M.V.A., C.R., E.A.M; Supervision: M.V.A., F.B., C.R., E.A.M. Resources: F.B., C.R., E.A.M.

