## Supplemental Figures for "Transposable elements drive regulatory and functional innovation of F-box genes"

### Supplemental Figures S1-S14.

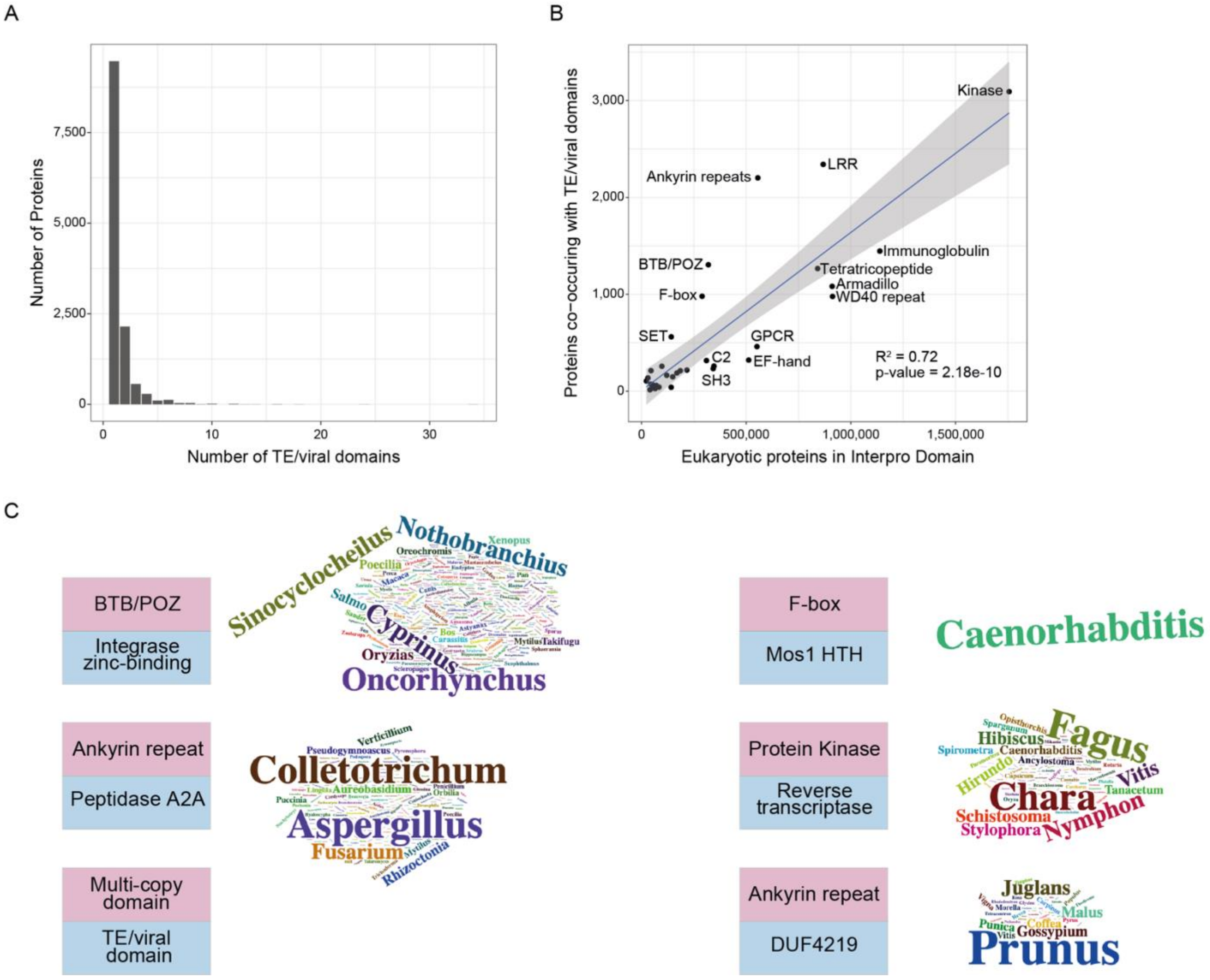


**Figure S1. Supplemental information on the eukaryotic proteins with multi-copy and TE- or virus-derived domains identified.** (**A**) Number of TE/viral domains per unique protein identified in the screen. (**B**) Association between the number of proteins in multi-copy domains and number of proteins with TE/viral domains. $R^2$ and p-value are indicated in the figure. (**C**) Word clouds representing the genera in which the proteins with these co-occurring domains are present. These are the largest sets of proteins with the same co-occurring domains, as shown in Figure 1E.

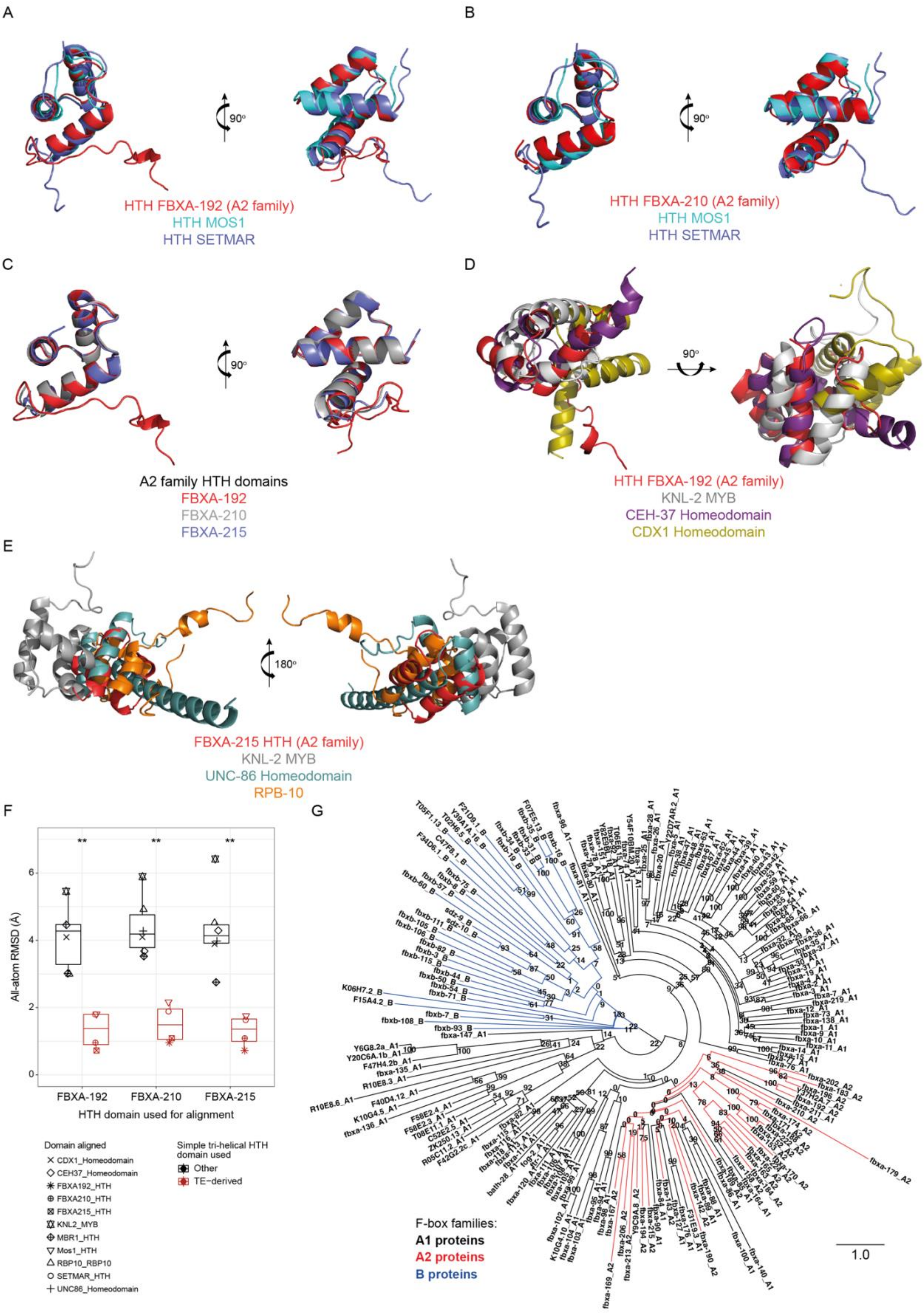
A
90°
HTH FBXA-192 (A2 family)
HTH MOS1
HTH SETMAR
B
90°
HTH FBXA-210 (A2 family)
HTH MOS1
HTH SETMAR
C
90°
A2 family HTH domains
FBXA-192
FBXA-210
FBXA-215
D
90°
HTH FBXA-192 (A2 family)
KNL-2 MYB
CEH-37 Homeodomain
CDX1 Homeodomain
E
180°
FBXA-215 HTH (A2 family)
KNL-2 MYB
UNC-86 Homeodomain
RPB-10
F
All-atom RMSD (Å)
FBXA-192
FBXA-210
FBXA-215
HTH domain used for alignment
Domain aligned
CDX1_Homeodomain
CEH37_Homeodomain
FBXA192_HTH
FBXA210_HTH
FBXA215_HTH
KNL2_MYB
MBR1_HTH
Mos1_HTH
RBP10_RBP10
SETMAR_HTH
UNC86_Homeodomain
Simple tri-helical HTH domain used
Other
TE-derived
G
F-box families:
A1 proteins
A2 proteins
B proteins
1.0

**Figure S2.**

**Figure S2. HTH domains of A2 family F-box genes are TE-derived and likely originate from a single TE capture.** (**A-B**) Structural alignments of the HTH domains of A2 family proteins, FBXA-192 in (A) and FBXA-210 in (B), with the HTH domains of *Drosophila mauritiana*'s Mos1 (PDB: 3HOT) and *Homo sapiens* SETMAR (PDB: 7S03). All these HTH domains are of the simple tri-helical type. (**C**) Structural alignment between the HTH domains of three A2 family proteins, demonstrating a consistent fold. (**D-E**) Structural alignments of HTH domains of A2 family proteins, FBXA-192 in (D) and FBXA-215 in (E), with other simple tri-helical HTH domains. (**F**) All-atom root mean square deviation (RMSD) of structural alignments between A2 family HTH domains with simple tri-helical HTH domains, both TE-derived and otherwise. Overall, A2 family HTH domains align consistently better with TE-derived HTH domains. P-values were computed with Wilcoxon rank-sum tests comparing RMSD values of alignments with TE-derived HTH domains and alignments with other simple tri-helical HTH domains (groups coloured in red versus black). (**G**) Maximum-likelihood phylogenetic tree of *C. elegans* F-box genes, as in Figure 2D but with branch lengths. The legend shows the colour code for all three gene families. Numbers at internal branches indicate bootstrap support values (100 pseudoreplicates).

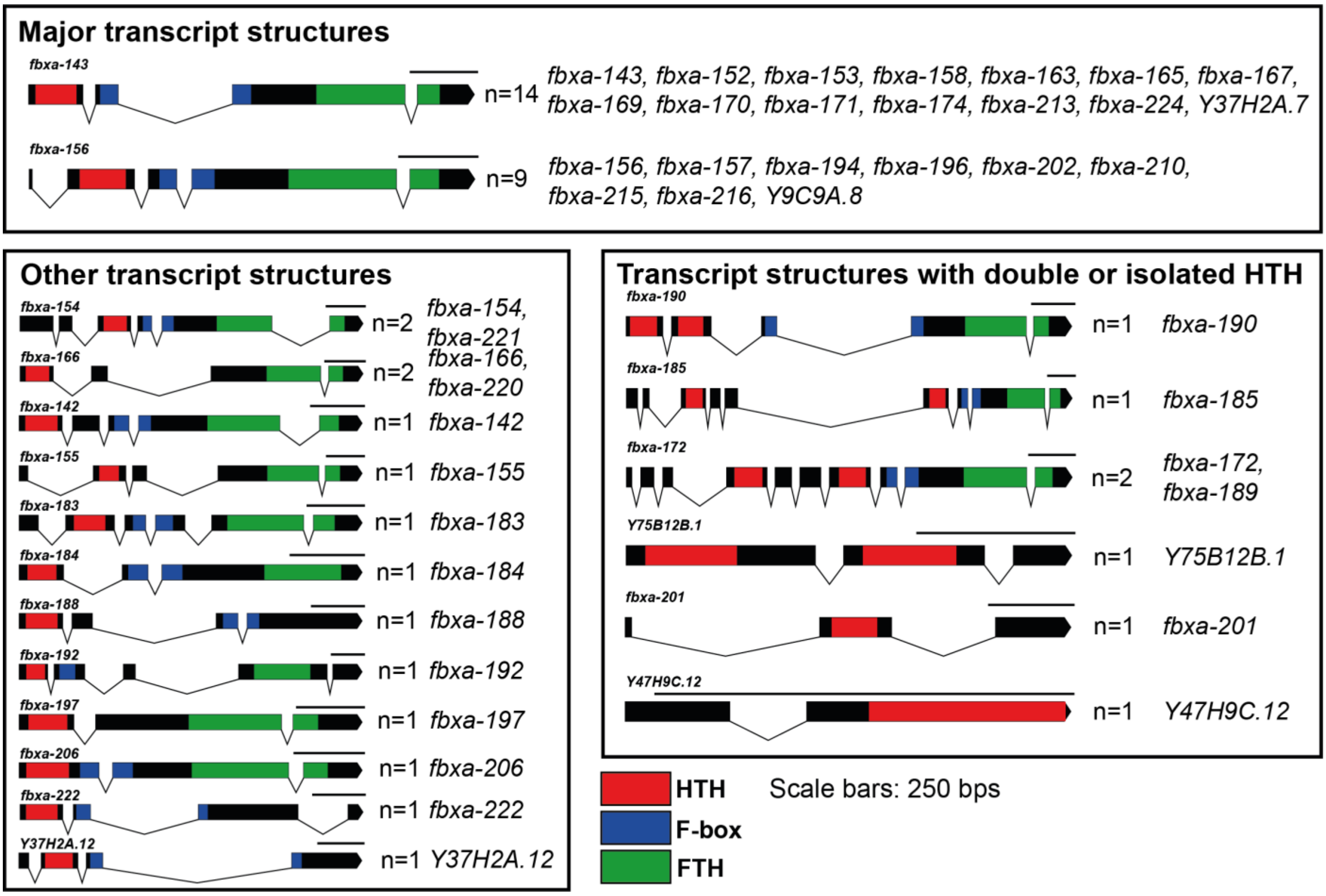


**Figure S3. F-box A2 genes have a defined architecture.** Transcript structure of all genes encoding an HTH domain in the *C. elegans* genome. Different exons are shown as black bars, and annotated domains are colour-coded. HTH domains are always encoded within a single exon and are always N-terminal relative to F-box and FTH domains. Some transcripts have two HTH domains of the same type (IPR041426). The representative scaled transcript structures were produced using an online resource (http://www.wormweb.org/exonintron). FTH, FOG-2 homology domain; HTH, Helix-turn-helix.

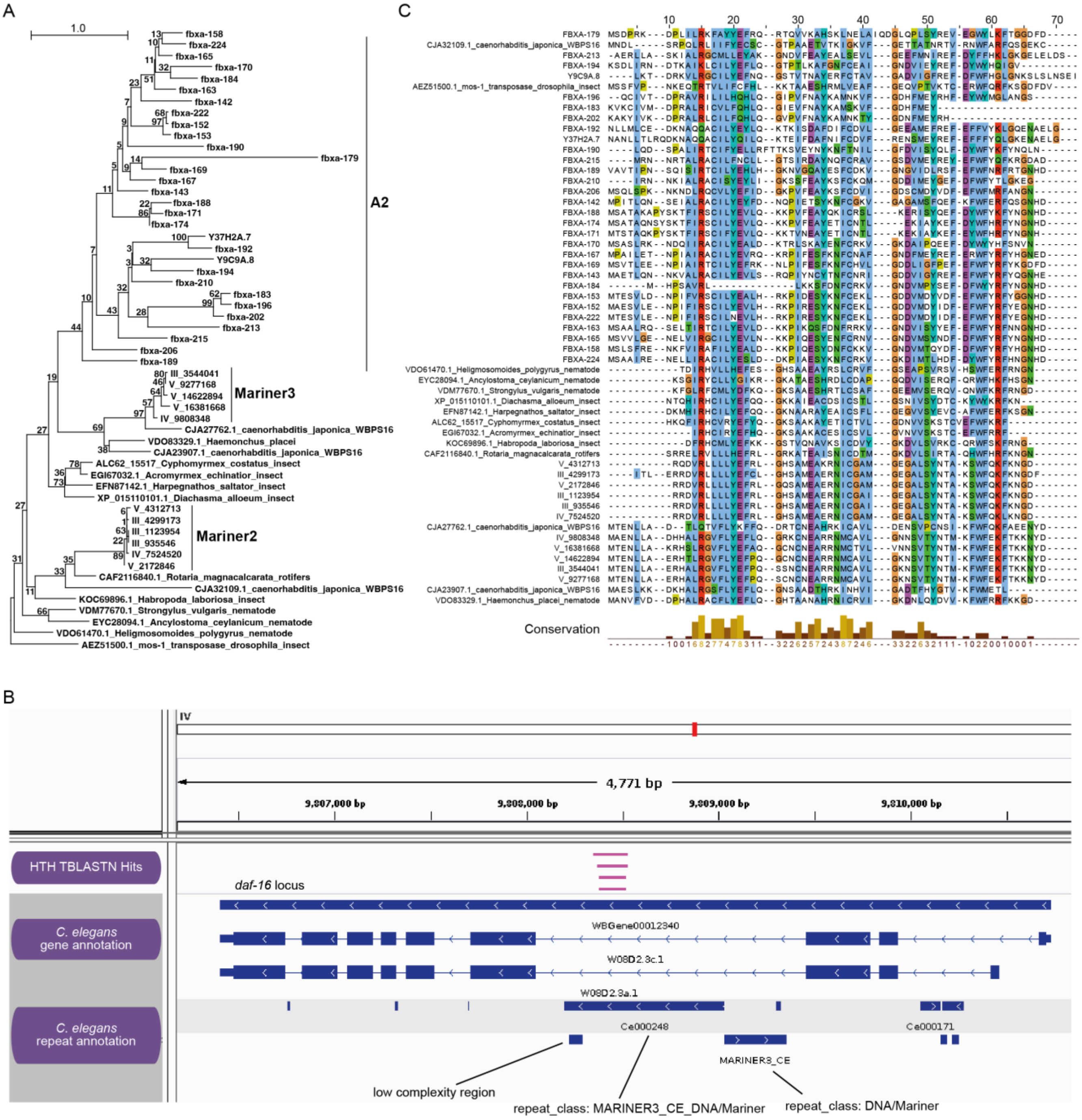
A
1.0
fbxa-158
fbxa-224
fbxa-165
fbxa-170
fbxa-184
fbxa-163
fbxa-142
fbxa-222
fbxa-152
fbxa-153
fbxa-190
fbxa-179
fbxa-169
fbxa-167
fbxa-143
fbxa-188
fbxa-171
fbxa-174
Y37H2A.7
fbxa-192
Y9C9A.8
fbxa-194
fbxa-210
fbxa-183
fbxa-196
fbxa-202
fbxa-213
fbxa-215
fbxa-206
fbxa-189
A2
III_3544041
V_9277168
V_14622894
V_16381668
IV_9808348
Mariner3
CJA27762.1_caenorhabditis_japonica_WBPS16
VDO83329.1_Haemonchus_placei
CJA23907.1_caenorhabditis_japonica_WBPS16
ALC62_15517_Cyphomyrmex_costatus_insect
EGI67032.1_Acromyrmex_echinatior_insect
EFN87142.1_Harpegnathos_saltator_insect
XP_015110101.1_Diachasma_alloeum_insect
V_4312713
III_4299173
III_1123954
III_935546
IV_7524520
V_2172846
Mariner2
CAF2116840.1_Rotaria_magnacalcarata_rotifers
CJA32109.1_caenorhabditis_japonica_WBPS16
KOC69896.1_Habropoda_laboriosa_insect
VDM77670.1_Strongylus_vulgaris_nematode
EYC28094.1_Ancylostoma_ceylanicum_nematode
VDO61470.1_Heligmosomoides_polygyrus_nematode
AEZ51500.1_mos-1_transposase_drosophila_insect
C
Conservation
B
IV
4,771 bp
9,807,000 bp
9,808,000 bp
9,809,000 bp
9,810,000 bp
HTH TBLASTN Hits
daf-16 locus
C. elegans gene annotation
WBGene00012340
W08D2.3c.1
W08D2.3a.1
C. elegans repeat annotation
Ce000248
Ce000171
MARINER3_CE
low complexity region
repeat_class: MARINER3_CE_DNA/Mariner
repeat_class: DNA/Mariner

**Figure S4.**

**Figure S4. The HTH domain of F-box A2 family factors originated in a Mariner TE**. (**A**) The likely donor of the HTH domain was an endemic TE. Phylogenetic analysis was performed for the HTH domains from F-box genes of the A2 family, their most closely-related sequences outside the Elegans group, and non-coding hits in the *C. elegans* genome. One of the most closely related sequences corresponds to an intronic Mariner element in *C. elegans*, see (B) and (C), which suggests that the HTH domain was derived from an endemic TE. (**B**) Genomic locus of the Mariner element in *C. elegans* that represents the closest relative to the HTH domains in F-box genes of the A2 family. Gene annotations and genomic coordinates of TBLASTN hits were visualized in the Integrative Genomics Viewer (Robinson et al. 2011). The screenshot shows a 4.8 kb region spanning the *daf-16* locus. The *C. elegans* gene *daf-16* has two isoforms. The largest intron contains sequences that derive from a Mariner transposon. (**C**) Multiple sequence alignment of HTH domains from diverse taxa generated using the MUSCLE tool and visualized with Jalview v2.11.4 (Waterhouse et al. 2009). Amino acid residues were coloured according to the colour scheme of Clustal X.

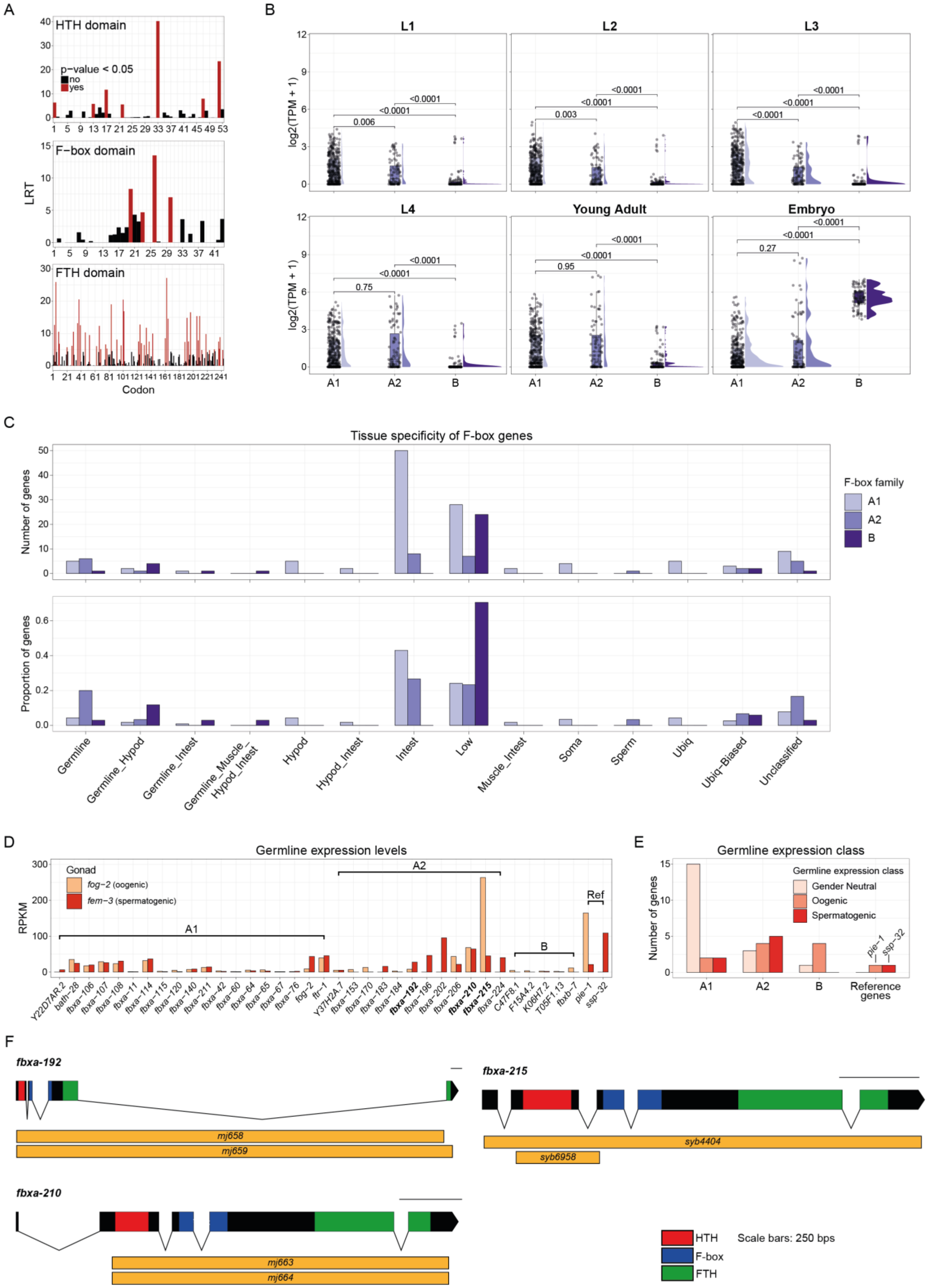
A
HTH domain
p−value < 0.05
no
yes
F−box domain
FTH domain
LRT
Codon
B
L1
L2
L3
L4
Young Adult
Embryo
log2(TPM + 1)
A1
A2
B
C
Tissue specificity of F-box genes
Number of genes
Proportion of genes
F-box family
Germline
Germline_Hypod
Germline_Intest
Germline_Muscle_Hypod_Intest
Hypod
Hypod_Intest
Intest
Low
Muscle_Intest
Soma
Sperm
Ubiq
Ubiq−Biased
Unclassified
D
Germline expression levels
RPKM
Gonad
fog−2 (oogenic)
fem−3 (spermatogenic)
Ref
E
Germline expression class
Number of genes
Gender Neutral
Oogenic
Spermatogenic
Reference genes
pie−1
ssp−32
F
fbxa-192
fbxa-215
fbxa-210
mj658
mj659
syb4404
syb6958
mj663
mj664
HTH
F-box
FTH
Scale bars: 250 bps

**Figure S5.**

**Figure S5. Selection and expression of nematode F-box factors.** (**A**) Panel shows the likelihood of codons to be under positive selection according to Mixed Effects Model of Evolution (MEME) (Murrell et al. 2012) across the alignment of all F-box A2 coding sequences in the *Caenorhabditis* genus. The C-terminal FTH domain of these proteins has a higher number of codons very likely under positive selection, when compared to the HTH and F-box domains (codons highlighted in red). LRT, likelihood ratio test. (**B**) Expression of F-box genes (split according to their family, either A1, A2, or B) throughout *C. elegans* development (larval stages L1, L2, L3, L4, young adult worms and mixed-staged embryos). Expression in log2-transformed transcripts per million (TPM). P-values were calculated with Wilcoxon rank-sum tests (using Benjamini & Hochberg correction). Publicly available expression data was used (Almeida et al. 2019). (**C**) Tissue specificity of F-box gene expression in *C. elegans* (split according to their family, either A1, A2, or B), as classified by another study (Serizay et al. 2020). Panel above shows the absolute number of genes in each category, while the panel below represents the proportion of genes of each family that is assigned to a particular category. (**D**) Expression data of F-box genes in *C. elegans* gonads isolated from female (oogenic, with *fog-2* mutation) and male (spermatogenic, with *fem-3* mutation) animals, using publicly available data (Ortiz et al. 2014). Only F-box genes with detectable expression in the germline are shown. Bold highlighting denotes three of the most highly expressed F-box A2 genes in the germline, which were mutated and followed up functionally. Two reference genes known to be enriched in oogenic or spermatogenic gonads are included: *pie-1* and *ssp-32*, respectively (Ortiz et al. 2014). (**E**) Number of F-box genes of each family classified as having gender neutral, oogenic, or spermatogenic germline expression, according to published datasets (Ortiz et al. 2014). (**F**) Schematics depicting the mutant alleles generated in this study. Range of deletions are indicated as orange boxes, along with the deletion allele name.

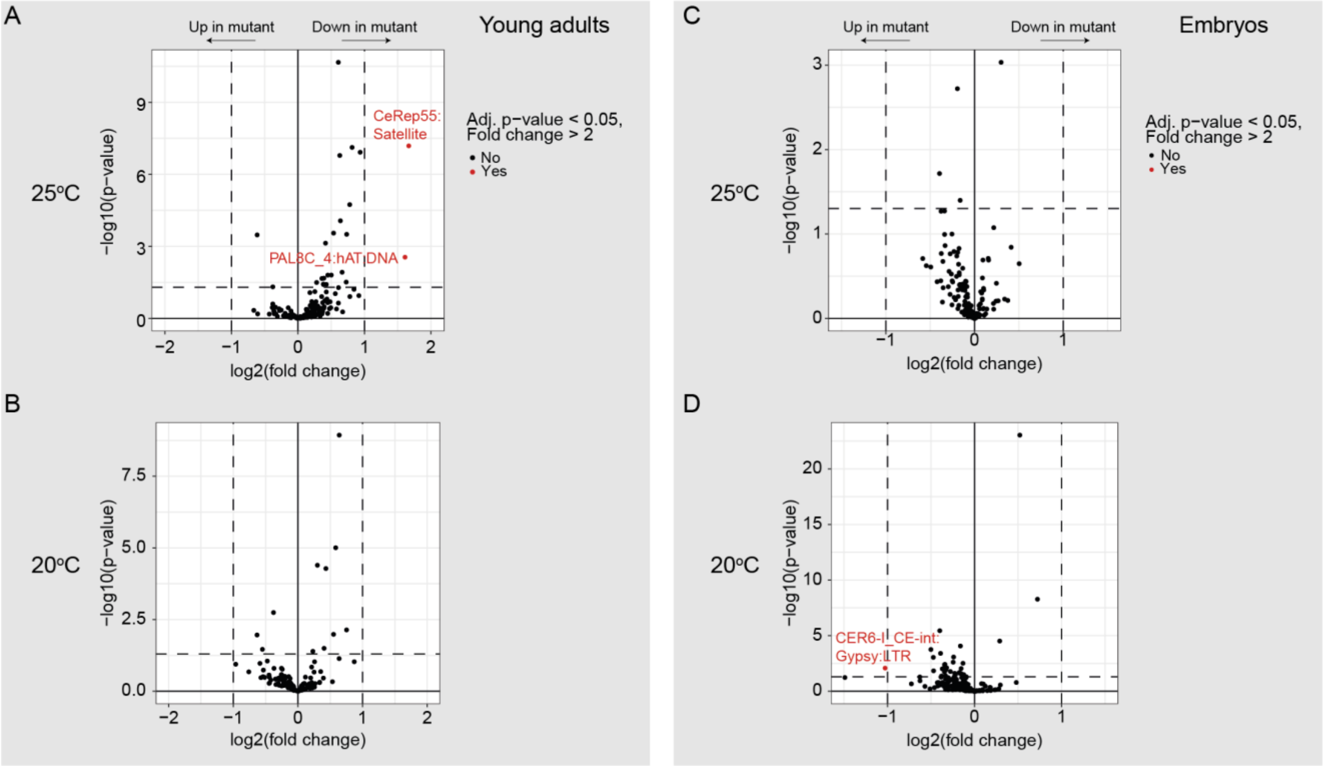


**Figure S6. FBXA-215 is unlikely to have a consistent role in transcriptional regulation of TEs.** (**A-D**) mRNA-sequencing results showing differential TE family expression in wild-type N2 and *fbxa-215* mutant animals. Using young adult animals (A-B) and mixed-stage embryos (C-D), grown at 20ºC (B, D) or 25ºC (A, C). Differentially expressed TE families are labelled in red in the volcano plots (defined by adjusted p-value < 0.05 and fold change > 2).

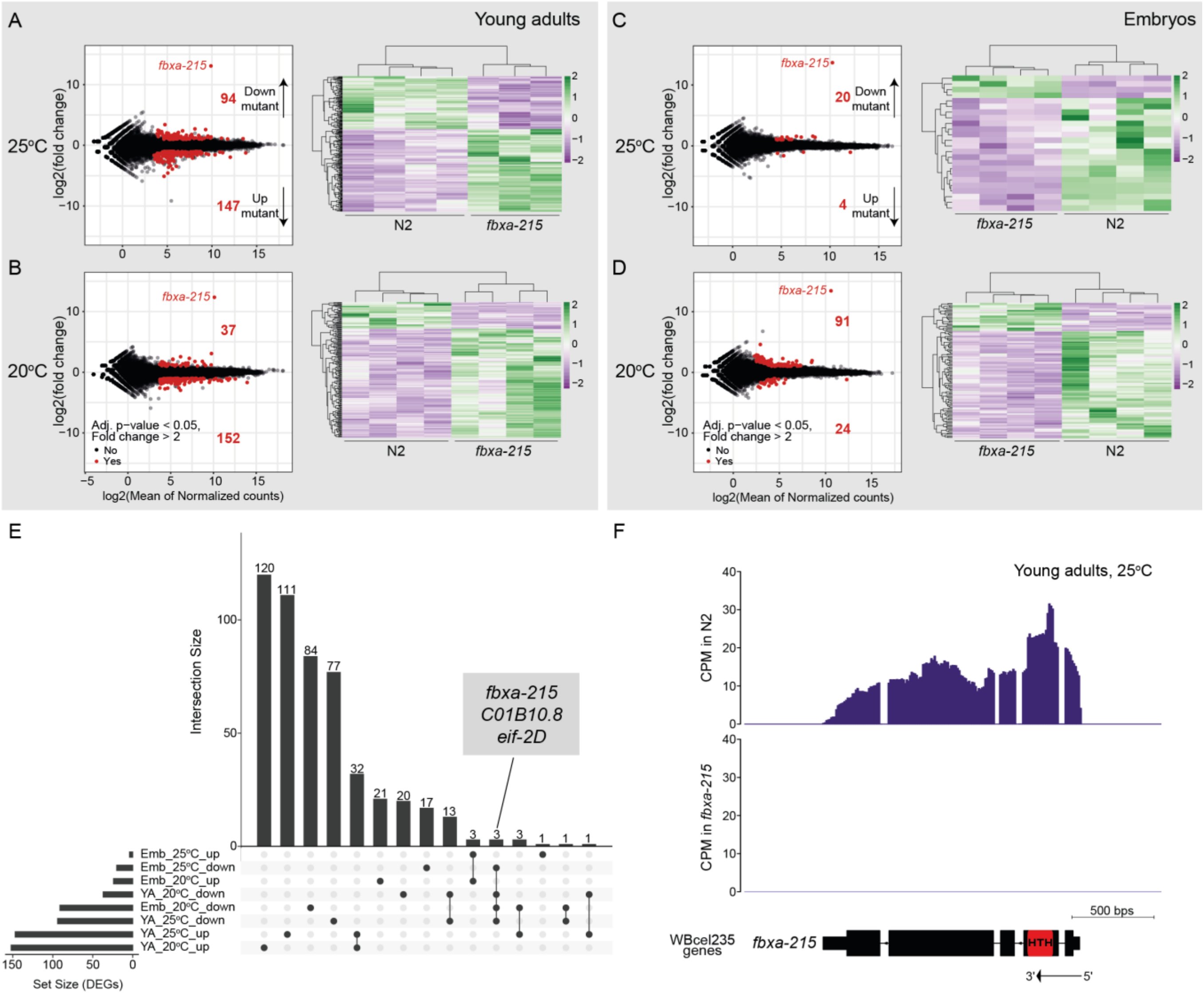
A
Young adults
25ºC
fbxa-215
94 Down mutant
147 Up mutant
log2(fold change)
N2
fbxa-215
B
20ºC
37
152
Adj. p-value < 0.05, Fold change > 2
No
Yes
log2(Mean of Normalized counts)
C
Embryos
20 Down mutant
4 Up mutant
D
91
24
E
Intersection Size
120
111
84
77
32
21
20
17
13
3
3
3
1
1
1
fbxa-215
C01B10.8
eif-2D
Emb_25ºC_up
Emb_25ºC_down
Emb_20ºC_up
YA_20ºC_down
Emb_20ºC_down
YA_25ºC_down
YA_25ºC_up
YA_20ºC_up
Set Size (DEGs)
F
Young adults, 25ºC
CPM in N2
CPM in fbxa-215
500 bps
WBcel235 genes
fbxa-215
HTH
3'
5'

**Figure S7.**

**Figure S7. Unlikely role for FBXA-215 in transcriptional gene regulation.** (**A-F**) mRNA-sequencing results showing differentially expressed genes between wild-type N2 and *fbxa-215* mutant worms. Using young adult animals (A-B) and mixed-stage embryos (C-D), grown at 20$^{o}$C (B, D) or 25$^{o}$C (A, C). In (A-D), the left panels are MA plots, and the right panels are heatmaps (with expression normalised using z-score), including only the differentially expressed genes (labelled in red in the MA plots, differential expression defined as follows: adjusted p-value < 0.05 and fold change > 2). *fbxa-215*, the mutated gene, is downregulated in all growth conditions and stages, and its datapoint is labelled in the MA plots in (A-D). (**E**) UpSet plot with the differentially expressed genes shared by different developmental stages and growth conditions. Three genes that are downregulated in mutants in all growth conditions and stages are indicated, and include *fbxa-215*, the mutated gene. (**F**) Genome tracks showing expression of the *fbxa-215* locus, as Counts per Million (CPM). Reassuringly, *fbxa-215* mutants, lacking the entire coding sequence of *fbxa-215*, have no detectable expression. The HTH domain is highlighted in red. These genome tracks represent data from young adults grown at 25$^{o}$C. These results are representative and identical to the genome tracks of this locus in other growth conditions and stages, corroborating the strong downregulation of *fbxa-215* observed in (A-D).

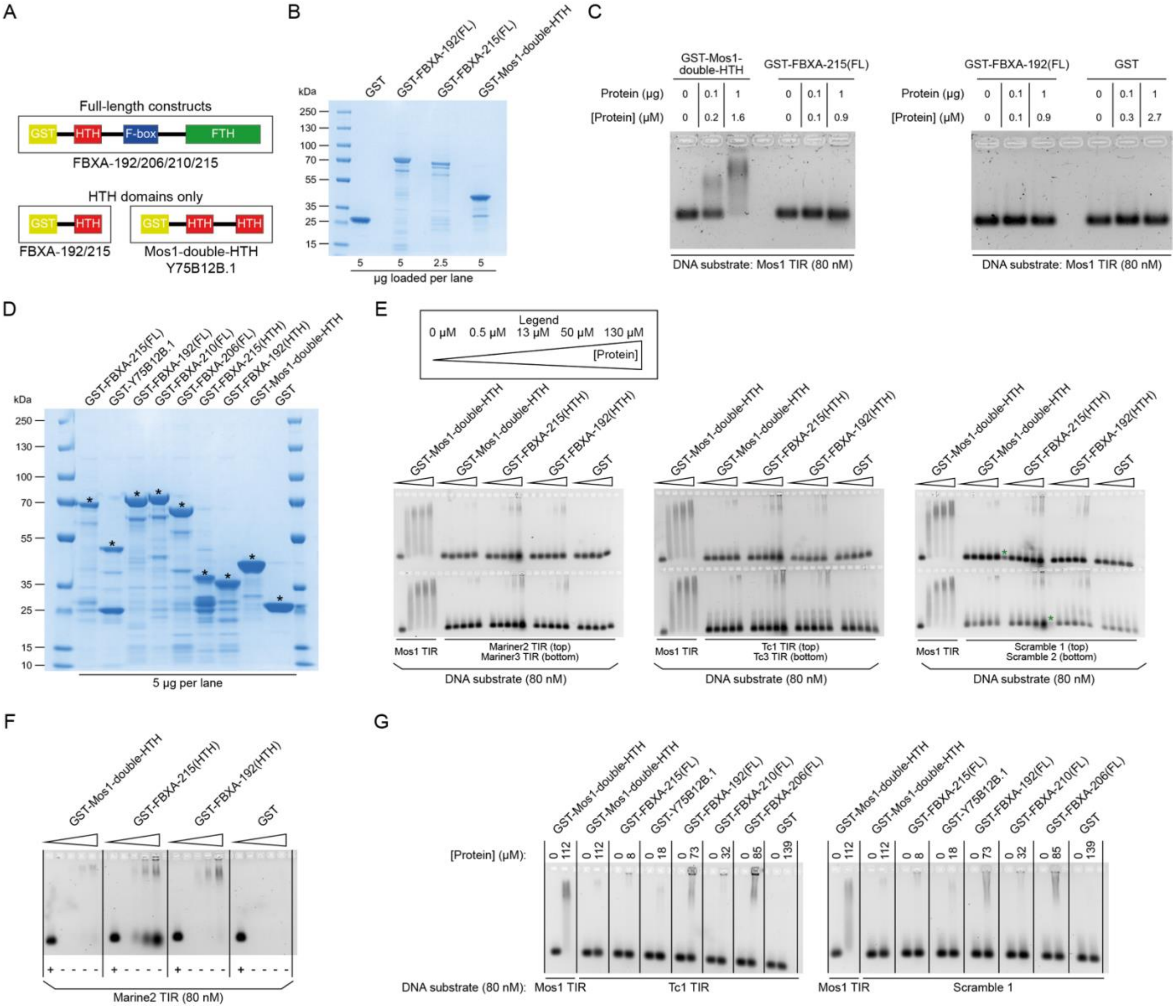
A
Full-length constructs
GST
HTH
F-box
FTH
FBXA-192/206/210/215
HTH domains only
FBXA-192/215
Mos1-double-HTH
Y75B12B.1
B
GST
GST-FBXA-192(FL)
GST-FBXA-215(FL)
GST-Mos1-double-HTH
kDa
250
130
100
70
55
35
25
15
5 5 2.5 5
μg loaded per lane
C
GST-Mos1-double-HTH
GST-FBXA-215(FL)
GST-FBXA-192(FL)
GST
Protein (μg)
[Protein] (μM)
DNA substrate: Mos1 TIR (80 nM)
D
GST-FBXA-215(FL)
GST-Y75B12B.1
GST-FBXA-192(FL)
GST-FBXA-210(FL)
GST-FBXA-206(FL)
GST-FBXA-215(HTH)
GST-FBXA-192(HTH)
GST-Mos1-double-HTH
GST
kDa
5 μg per lane
E
Legend
0 μM 0.5 μM 13 μM 50 μM 130 μM
[Protein]
Mos1 TIR
Mariner2 TIR (top)
Mariner3 TIR (bottom)
Tc1 TIR (top)
Tc3 TIR (bottom)
Scramble 1 (top)
Scramble 2 (bottom)
DNA substrate (80 nM)
F
GST-Mos1-double-HTH
GST-FBXA-215(HTH)
GST-FBXA-192(HTH)
GST
Marine2 TIR (80 nM)
G
GST-Mos1-double-HTH
GST-FBXA-215(FL)
GST-Y75B12B.1
GST-FBXA-192(FL)
GST-FBXA-210(FL)
GST-FBXA-206(FL)
GST
[Protein] (μM):
DNA substrate (80 nM):
Mos1 TIR
Tc1 TIR
Scramble 1

**Figure S8.**

**Figure S8. F-box A2 proteins and their HTH domains do not show appreciable binding to Tc1/mariner elements *in vitro*.** (**A**) Schematics showing the domain structures of the proteins expressed in *Escherichia coli* and purified to conduct electrophoretic mobility shift assays (EMSAs) to test DNA-binding. The proteins and domains of interest were expressed together with an N-terminal Glutathione S-transferase (GST) tag to increase solubility. We cloned a set of GST-tagged full-length F-box A2 proteins, as well as versions including only the HTH domains of FBXA-192 and FBXA-215. As a control we included the two HTH domains of the Mos1 transposase, which are related to the F-box A2 HTH domains and are known to bind to DNA (Richardson et al. 2009). In addition, we included the full length Y75B12B.1, a protein that is not an F-box A2 protein, but is predicted to encode two HTH domains of the same family as the HTH domains of F-box A2 proteins (see Figure S3). This protein was included to understand if two HTH domains are required for DNA-binding. (**B**) SDS-PAGE gel stained with Coomassie InstantBlue, showing purified GST fusion proteins for a pilot EMSA experiment. (**C**) EMSA pilot experiment showing high-affinity binding of the isolated Mos1 transposase double-HTH DNA-binding domain to its cognate terminal inverted repeat (TIR). Experiment was repeated three times. Full-length (FL) FBXA-192 and FBXA-215 are not able to bind Mos1 TIR. (**D**) SDS-PAGE gel stained with Coomassie InstantBlue showing the protein purifications for subsequent EMSA assays. Asterisks indicate expected sizes. (**E**) EMSA testing the binding of isolated HTH domains from FBXA-192 and FBXA-215 to the TIRs of eroded *C. elegans* Mariner2 and Mariner3 DNA TEs (left panel), to the TIRs of active *C. elegans* Tc1 and Tc3 DNA TEs (middle panel), and to two randomly generated DNA sequences (right panel). Gels are stained with SYBR Gold. Green asterisks indicate signal derived from spillage of adjacent wells when loading the gel. No significant binding is observed to any of the DNA substrates we introduced, with the exception of the positive control of Mos1 and its TIR. The faint slower migrating signal observed in higher concentrations of the protein is due to contaminating nucleic acid associated with the GST-purified constructs, see (F). (**F**) Examining protein preparations used in (E) on a native EMSA agarose gel without any DNA substrate in the binding reactions and staining with SYBR Gold. The first wells with no protein added were loaded with DNA as a control. Signal is observed on the higher protein concentrations and is thus not due to the DNA substrate we introduce, but instead very likely to be nucleic acids carried over from the protein purification in *E. coli*. This nucleic acid contaminant is observed in GST-Mos1-double-HTH constructs as well, and it does not affect binding of these proteins to the known DNA substrate. Thus, this contaminant is unlikely to preclude binding of F-box A2 proteins to bona fide DNA substrates. In (E-F), protein concentration range follows the legend in (E). (**G**) EMSA testing the binding of full-length HTH-domain-containing proteins to the TIR of the active *C. elegans* Tc1 DNA TE, and to a randomly generated DNA sequence. Protein concentrations were not standardised as in (E-F) and are indicated in the figure. (C, E-G) Molarities represent final concentrations in the binding reaction.

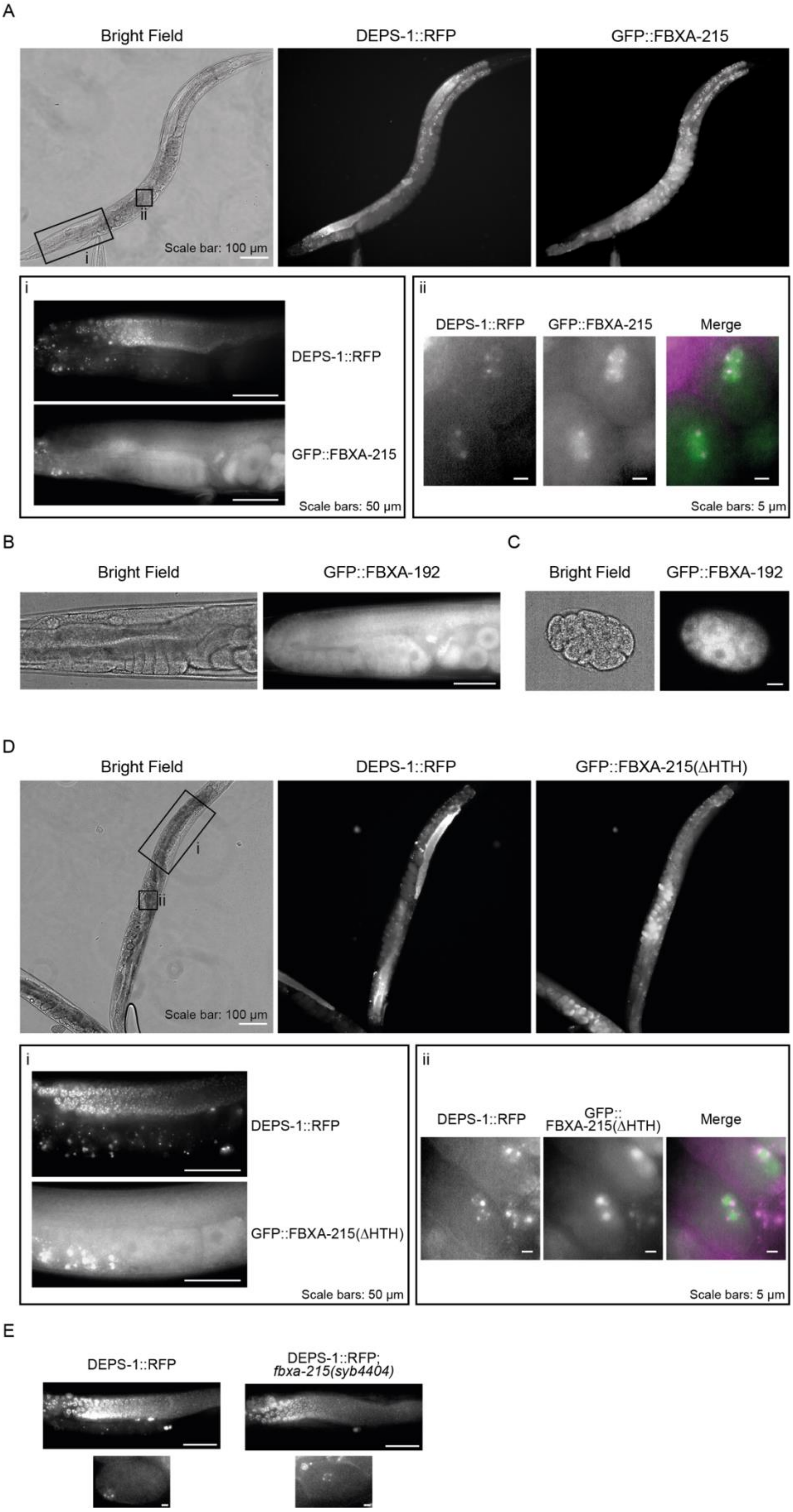
A
Bright Field
DEPS-1::RFP
GFP::FBXA-215
ii
i
Scale bar: 100 µm
i
DEPS-1::RFP
GFP::FBXA-215
Scale bars: 50 µm
ii
DEPS-1::RFP
GFP::FBXA-215
Merge
Scale bars: 5 µm
B
Bright Field
GFP::FBXA-192
C
Bright Field
GFP::FBXA-192
D
Bright Field
DEPS-1::RFP
GFP::FBXA-215(ΔHTH)
i
ii
Scale bar: 100 µm
i
DEPS-1::RFP
GFP::FBXA-215(ΔHTH)
Scale bars: 50 µm
ii
DEPS-1::RFP
GFP::
FBXA-215(ΔHTH)
Merge
Scale bars: 5 µm
E
DEPS-1::RFP
DEPS-1::RFP;
fbxa-215(syb4404)

**Figure S9.**

**Figure S9. Expression and localisation of FBXA-192 and FBXA-215 in *C. elegans*.** (**A**) Bright field and fluorescence photomicrographs showing a representative worm expressing DEPS-1::RFP and GFP::FBXA-215 (wild-type FBXA-215). Inset (i) shows expression in the adult worm germline, while inset (ii) shows expression in embryos. Scales are indicated in the figure. (**B-C**) Representative images of expression of GFP::FBXA-192 in the adult germline (B) and in embryos (C). Scale bars in (B) and (C) are equivalent to 50 and 10 μm, respectively. (**D**) Photomicrographs showing a representative worm expressing DEPS-1::RFP and GFP::FBXA-215(ΔHTH). Inset (i) shows expression in the adult germline, while inset (ii) shows expression in embryos. Scales are indicated in the figure. (**E**) Germ granules are not affected by mutation of *fbxa-215*, evaluated by DEPS-1::RFP expression. Wild-type (left panels) and *fbxa-215(syb4404)* worms (right panels), do not show obvious differences. Scale bars in upper and lower panels depict 50 and 5 μm, respectively. (A-E) All worms of the same genotype had identical expression (n >10).

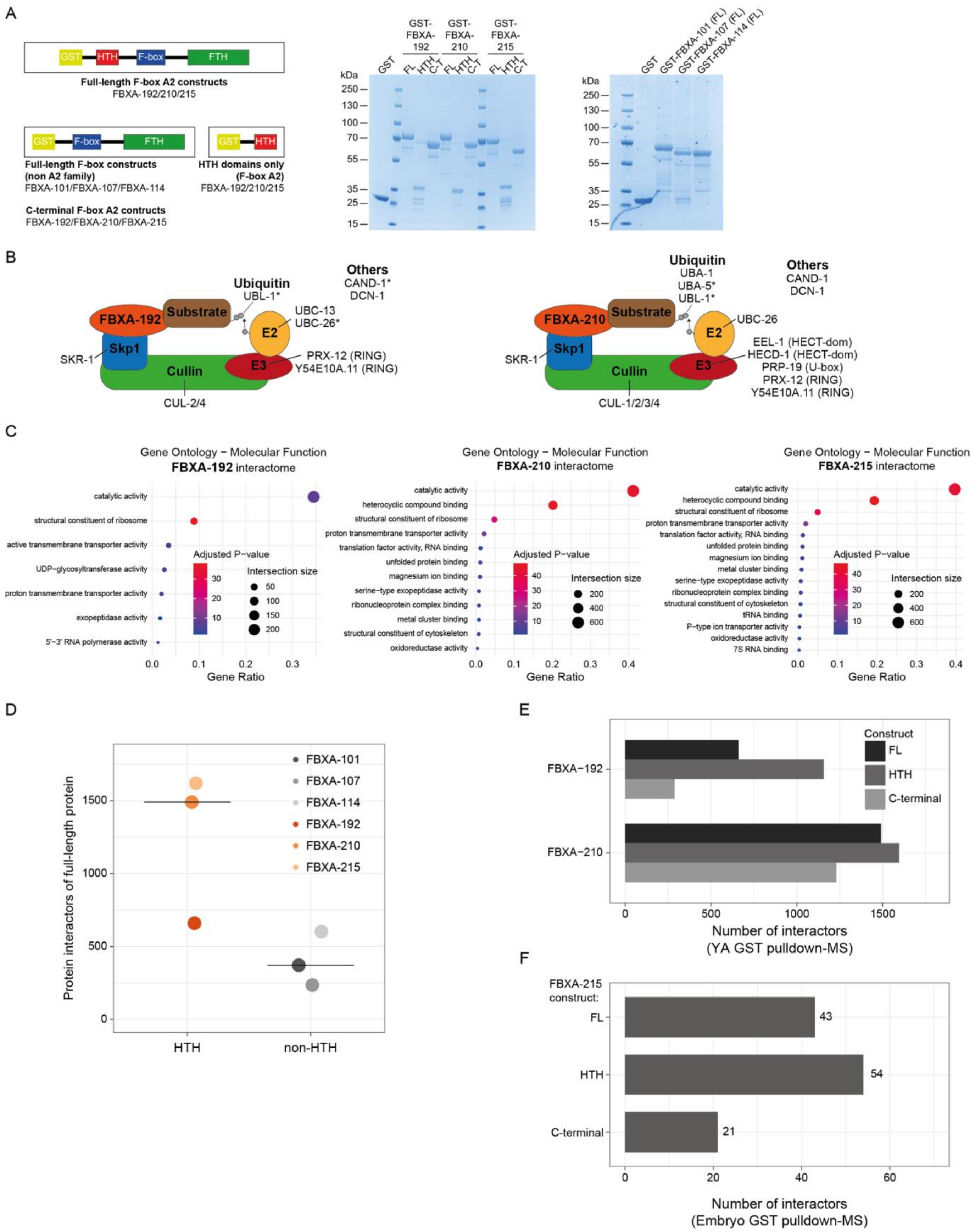
A
GST
HTH
F-box
FTH
Full-length F-box A2 constructs
FBXA-192/210/215
Full-length F-box constructs
(non A2 family)
FBXA-101/FBXA-107/FBXA-114
HTH domains only
(F-box A2)
FBXA-192/210/215
C-terminal F-box A2 contructs
FBXA-192/FBXA-210/FBXA-215
GST-FBXA-192
GST-FBXA-210
GST-FBXA-215
FL
HTH
C-T
kDa
250
130
100
70
55
35
25
15
GST-FBXA-101 (FL)
GST-FBXA-107 (FL)
GST-FBXA-114 (FL)
B
Ubiquitin
UBL-1*
Others
CAND-1*
DCN-1
FBXA-192
Substrate
Skp1
SKR-1
Cullin
CUL-2/4
E2
UBC-13
UBC-26*
E3
PRX-12 (RING)
Y54E10A.11 (RING)
Ubiquitin
UBA-1
UBA-5*
UBL-1*
Others
CAND-1
DCN-1
FBXA-210
UBC-26
EEL-1 (HECT-dom)
HECD-1 (HECT-dom)
PRP-19 (U-box)
PRX-12 (RING)
Y54E10A.11 (RING)
CUL-1/2/3/4
C
Gene Ontology − Molecular Function
FBXA-192 interactome
catalytic activity
structural constituent of ribosome
active transmembrane transporter activity
UDP-glycosyltransferase activity
proton transmembrane transporter activity
exopeptidase activity
5'-3' RNA polymerase activity
Adjusted P−value
Intersection size
Gene Ratio
Gene Ontology − Molecular Function
FBXA-210 interactome
heterocyclic compound binding
translation factor activity, RNA binding
unfolded protein binding
magnesium ion binding
serine−type exopeptidase activity
ribonucleoprotein complex binding
metal cluster binding
structural constituent of cytoskeleton
oxidoreductase activity
Gene Ontology − Molecular Function
FBXA-215 interactome
tRNA binding
P-type ion transporter activity
7S RNA binding
D
Protein interactors of full-length protein
FBXA-101
FBXA-107
FBXA-114
FBXA-192
FBXA-210
FBXA-215
HTH
non-HTH
E
Construct
FL
HTH
C-terminal
FBXA−192
FBXA−210
Number of interactors
(YA GST pulldown-MS)
F
FBXA-215
construct:
FL
HTH
C-terminal
43
54
21
Number of interactors
(Embryo GST pulldown-MS)

**Figure S10.**

**Figure S10. Defining the interactome of F-box A2 proteins.** (**A**) Left panel illustrates the domain structures of the Glutathione S-transferase (GST)-F-box fusion proteins constructed in this work. Middle and right panels show SDS-PAGE gels stained with Coomassie InstantBlue, with 5 μg GST-F-box fusion proteins purified from *Escherichia coli*. Middle panel shows F-box A2 proteins, while right panel shows F-box domain-containing proteins not belonging to the F-box A2 family. (**B**) Schematics representing the SCF complex factors detected in the interactome of FBXA-192 (left), and FBXA-210 (right). Asterisk indicates proteins that were classified as an interactor of the HTH domain, but not of the full-length protein. (**C**) Gene ontology (Molecular Function) of the interactome of F-box A2 proteins: FBXA-192 (left panel), FBXA-210 (middle panel), and FBXA-215 (right panel). (**D**) Number of interactors of F-box proteins, comparing F-box A2 proteins, which have an HTH domain, with other F-box domain-containing proteins lacking the TE-derived HTH domain. Horizontal bar represents the median. (**E**) Number of protein interactors of F-box A2 proteins defined by pulldown-mass spectrometry. Pulldowns were performed in young adult animal extracts using the full-length (FL), the HTH domain, and the C-terminal region of F-box A2 proteins. (**F**) Number of proteins interacting with full-length (FL) FBXA-215, its HTH domain, and C-terminal region, as determined by pulldown-mass spectrometry. Pulldowns were performed in embryo extracts. (B-F) See detailed results in Table S3.

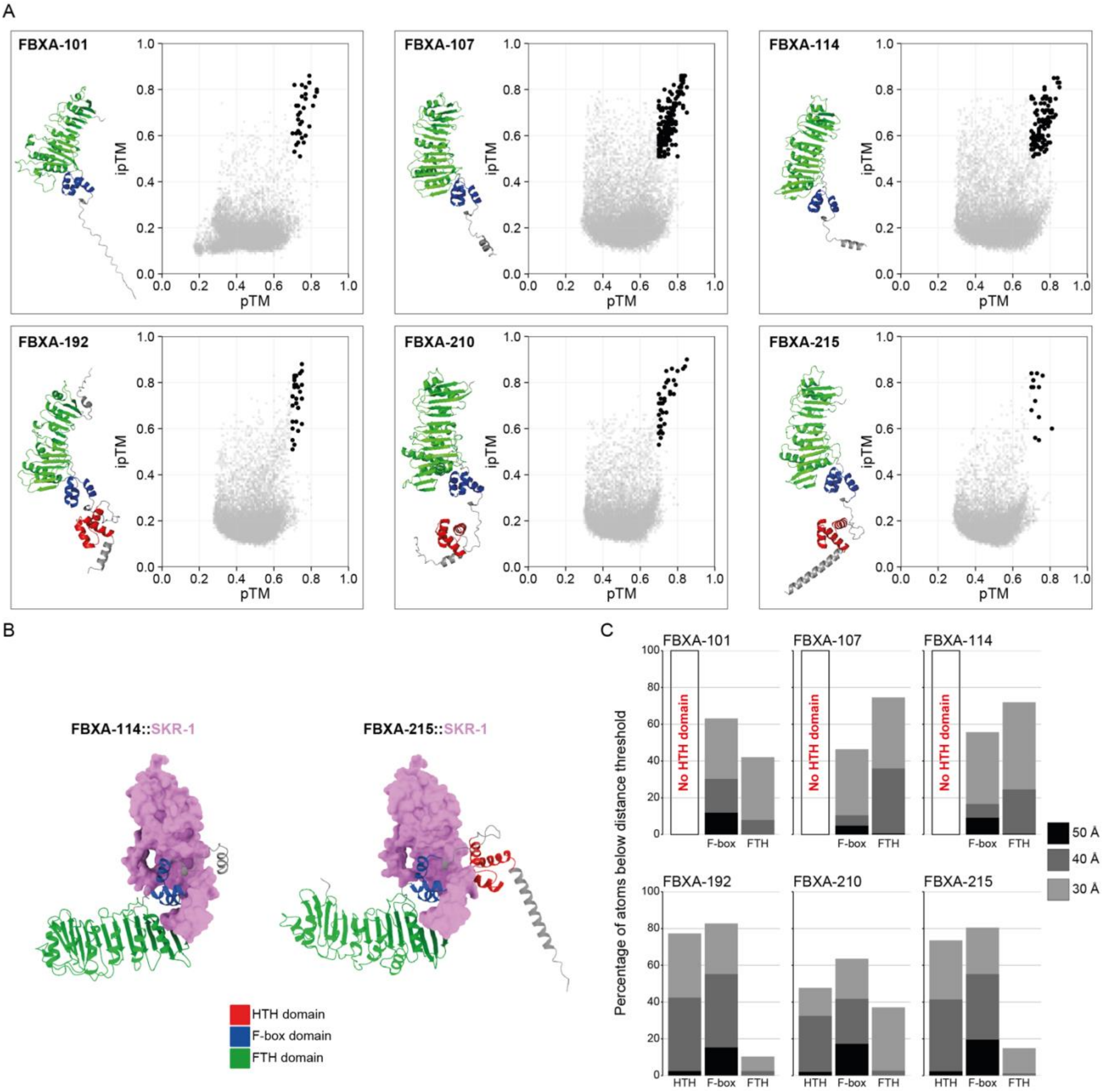
A
FBXA-101
FBXA-107
FBXA-114
FBXA-192
FBXA-210
FBXA-215
ipTM
pTM
B
FBXA-114::SKR-1
FBXA-215::SKR-1
HTH domain
F-box domain
FTH domain
C
Percentage of atoms below distance threshold
No HTH domain
HTH
F-box
FTH
50 Å
40 Å
30 Å

**Figure S11.**

**Figure S11. Additional data of the AlphaFold2 multimer screen for F-box interactors.** (**A**) AlphaFold2 multimer screen results for each F-box protein bait used. The pTM and ipTM values are shown for each interaction over the entire germline proteome of *C. elegans*. Black dots show the high-confidence interactions, defined by the highest pTM and ipTM (see Methods). AlphaFold2-modelled structures of the F-box protein baits are shown. (**B**) High-confidence interactions between representative F-box proteins and SKR-1, a Skp1 ortholog, modelled by AlphaFold2 multimer. The surface of SKR-1 is represented in pink. SKR-1 interfaces with the F-box domain, in blue, of FBXA-114 (not containing an HTH domain, on the left) and FBXA-215 (an HTH domain-containing F-box A2 protein, on the right). All F-box proteins used as baits in the screen, as represented in (A), have identical interactions with SKR-1, according to AlphaFold2. (A-B) colour key for the protein domains of the F-box protein structures is shown under (B). (**C**) Barplots showing the percentage of atoms of high-confidence interactors of F-box proteins, as determined in (A), below a certain threshold of distance to the F-box protein (30, 40, or 50 Å).

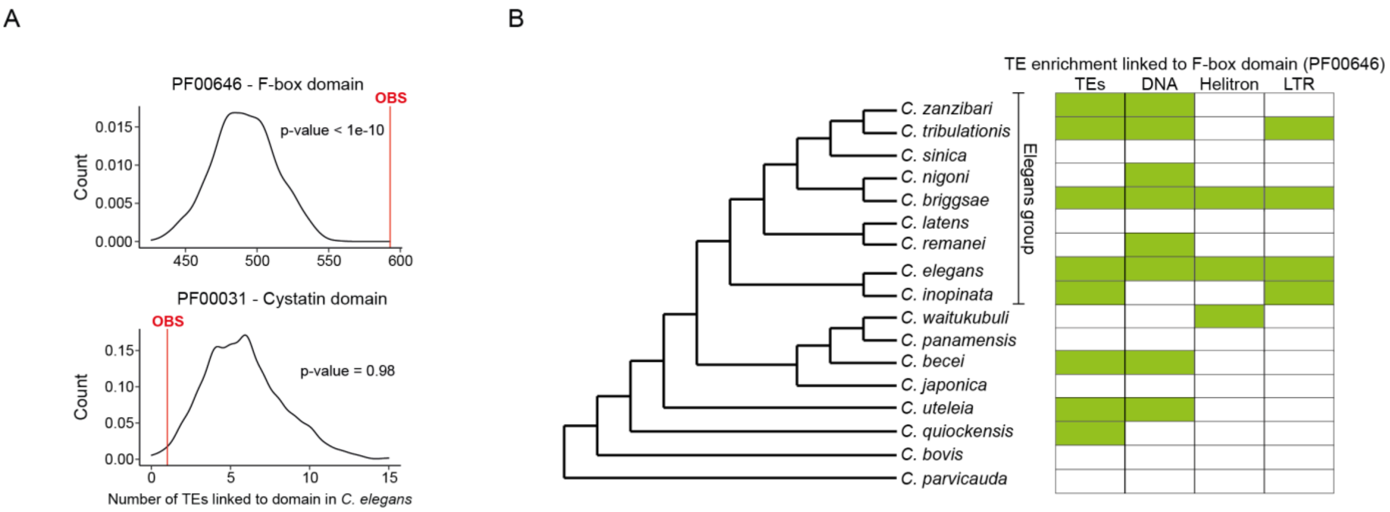


**Figure S12. F-box genes are located in TE-rich regions of *Caenorhabditis* genomes.** (**A**) Distribution of bootstrapped TE counts linked to all annotated protein-coding genes with a given protein domain. Here we show two example TEs distributions in *C. elegans* for genes with an annotated F-box domain and genes with a Cystatin domain, as a representative negative control. The observed, real number of TEs in the *C. elegans* genome is denoted with a red line. (**B**) Summary of TE enrichment linked to annotated protein-coding genes with an F-box domain, across 17 *Caenorhabditis* genomes. Statistically significant enrichment of TEs in F-box domain-containing genes is represented as green-coloured boxes (p-value < 0.05), whereas white-coloured boxes illustrate lack of significant TE enrichment (p-value ≥ 0.05). We tested for TE enrichment in general ("TEs" column) and for enrichment of specific TE classes.

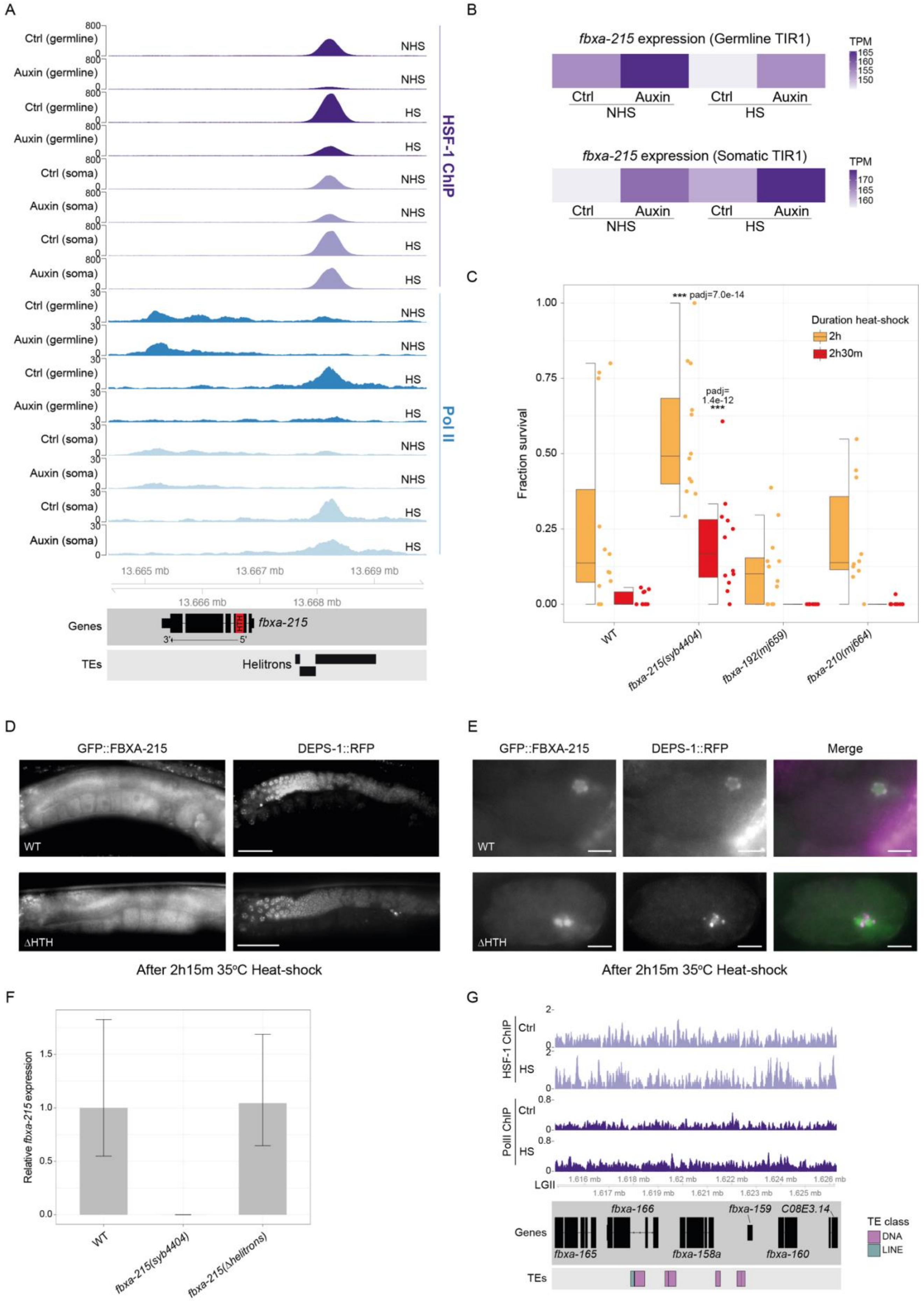
A
Ctrl (germline)
Auxin (germline)
Ctrl (soma)
Auxin (soma)
NHS
HS
HSF-1 ChIP
Pol II
13.665 mb
13.667 mb
13.669 mb
13.666 mb
13.668 mb
Genes
fbxa-215
TEs
Helitrons
B
fbxa-215 expression (Germline TIR1)
fbxa-215 expression (Somatic TIR1)
TPM
Ctrl
Auxin
C
*** padj=7.0e-14
padj=
1.4e-12
***
Duration heat-shock
2h
2h30m
Fraction survival
WT
fbxa-215(syb4404)
fbxa-192(mj659)
fbxa-210(mj664)
D
GFP::FBXA-215
DEPS-1::RFP
ΔHTH
After 2h15m 35°C Heat-shock
E
Merge
F
Relative fbxa-215 expression
fbxa-215(Δhelitrons)
G
LGII
fbxa-166
fbxa-159
C08E3.14
fbxa-165
fbxa-158a
fbxa-160
TE class
DNA
LINE
PolII ChIP

**Figure S13.**

**Figure S13. The molecular and phenotypic impact of integration of *fbxa-215* in a heat-shock response by HSF-1.** (**A**) Genome tracks centered at the *fbxa-215* locus, with HSF-1 (purple tracks) and RNA Polymerase II (blue tracks) ChIP-sequencing data, obtained from animals expressing TIR1 in the germline or in the soma. (**B**) Expression of *fbxa-215* in transcripts per million (TPM) in the same conditions as (A). (A-B) Publicly available ChIP-sequencing and RNA-sequencing data was used to produce these panels (Edwards et al. 2021). Ctrl, control with no auxin treatment; HS, Heat-shock; NHS, Non-heat-shocked. (**C**) Worm survival 24 hours after 37$^{o}$C heat-shock for 2h (orange) or 2h30m (red). Each figure represents two combined experiments. P-values show the results of Fisher's exact tests comparing survival of worm strains versus survival of wild-type (WT) N2. Horizontal lines in the boxes represent the median. The bottom and top of the box represent the 25$^{th}$ and 75$^{th}$ percentile. Whiskers include data points that are less than 1.5 x interquartile range away from the 25$^{th}$ and 75$^{th}$ percentile. (**D-E**) Fluorescence photomicrographs showing a representative adult worm (D) or embryo (E) expressing DEPS-1::RFP and GFP::FBXA-215 wild-type sequence (upper panels), or ΔHTH (lower panels), after 2h15m heat-shock at 35$^{o}$C. Compare with Figures 3A and S9 for localisation at a non-stressful temperature. All worms of the same genotype showed identical expression (n >10). Scale bars in (D) and (E) are equivalent to 50 and 10 $\mu$m, respectively. (**F**) RT-qPCR results showing *fbxa-215* expression relative to *pmp-3* in the indicated strains. Animals used for this experiment were grown at a non-stressful temperature (20$^{o}$C). (**G**) Genome tracks showing HSF-1 and RNA Polymerase II ChIP-sequencing data from young adult worms (Li et al. 2016) at the *fbxa-158* locus. Ctrl, Control; HS, Heat-shock. Tracks show a genome region centered at *fbxa-158* +/- 5 Kb. No Helitron TEs are annotated in this region in the wild-type N2 genome reference.

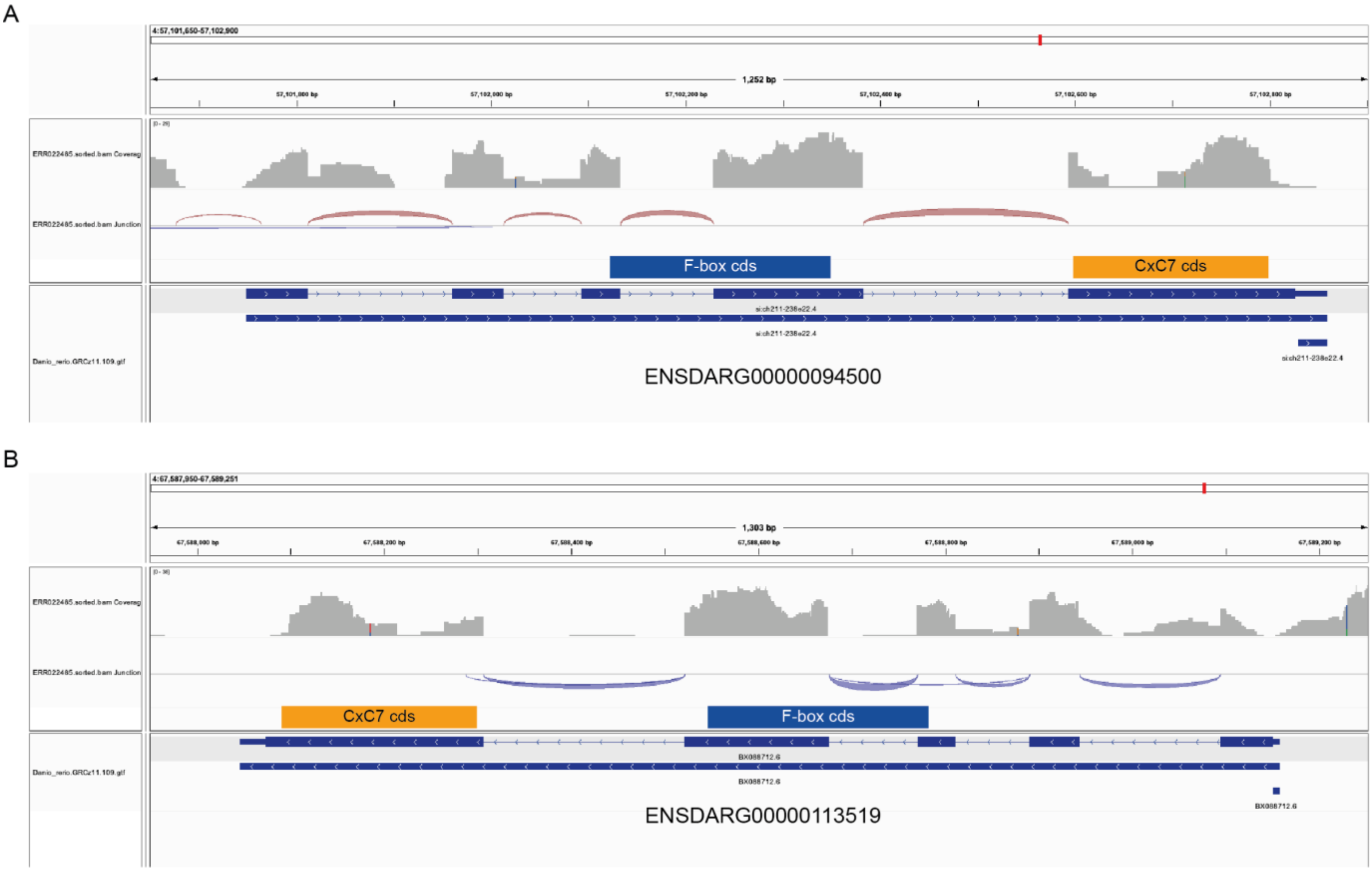


**Figure S14. F-box::CxC7 genes are expressed as a single transcriptional unit.** (**A-B**) Integrative Genomics Viewer (Robinson et al. 2011) screenshots showing publicly available RNA-sequencing data (Collins et al. 2012) mapped to two F-box::CxC7 genes, ENSDARG00000094500 (A) and ENSDARG00000113519 (B). The boxes above the gene annotation tracks denote the coding sequences (cds) of the genes encoding the F-box (blue) and CxC7 (orange) protein domains. The upper tracks have information on coverage and on exon-exon junctions. The latter is defined by supporting reads spanning the exon-exon junction. The junction tracks support that the annotated exons coding the F-box protein domain are spliced together with the exon containing the CxC7 domain.
